# Temperature-dependent biofilm and sublancin production immobilise arsenic and antibiotic resistance gene mobility in soil system

**DOI:** 10.64898/2026.01.11.698866

**Authors:** Arnab Majumdar, Ioly Kotta-Loizou, Martin Buck, Tarit Roychowdhury

**Author notes:** Correspondence- Dr. Arnab Majumdar, Ph.D., MRSB, AMRSC (,).

## Abstract

Climate change-induced warming and arsenic soil contamination synergistically threaten agricultural sustainability by restructuring microbial communities and accelerating antibiotic resistance evolution. Here, we demonstrate that Bacillus subtilis 168-derived sublancin, a temperature-optimised glycosylated antimicrobial peptide, simultaneously maximises microbial arsenic sequestration and suppresses antibiotic resistance gene (ARG) horizontal transfer. Temperature-dependent biofilm formation increased progressively from 25–35°C (absorbance: 0.37–1.2), correlating with enhanced arsenic accumulation within the extracellular polymeric substance matrix (74% increase in weight percentage). Time-of-flight secondary ion mass spectrometry confirmed maximum arsenic sequestration at 35°C (total counts: 4.725×10⁴). Sublancin production peaked at 30.8°C (129.72 mg/L) and significantly reduced soil microbial biomass carbon by 33% (p < 0.001), with selective suppression of Gram-positive bacteria (42%–28% survival reduction). Conjugation assays revealed sublancin bio-amendment dramatically suppressed horizontal ARG transfer, reducing HGT frequency by 74.7% (6.64 × 10⁻³ to 1.68 × 10⁻³, p < 0.001) through dual mechanisms: decreased donor viability and reduced conjugation efficiency. Gene ontology enrichment identified coordinated upregulation of exopolysaccharide synthesis (FDR ~1.0×10⁻²⁷), phosphotransferase systems, and arsenic resistance pathways, alongside downregulation of conventional antibiotic resistance genes. These findings establish sublancin as a dual-function soil amendment simultaneously addressing two critical agricultural threats: heavy metal bioaccumulation and antibiotic resistance dissemination under climate warming.

## Introduction

Climate change-induced warming, advancing at 1.1°C since 1850-1900 with projections reaching 1.5°C within decades (NASA, 2024), fundamentally threatens agricultural soil systems through interconnected mechanisms. Warming accelerates soil organic matter decomposition, triggering nutrient loss and reduced soil quality, whilst simultaneously restructuring soil microbial communities, bacterial and fungal diversity declining by 16.0% and 19.7% respectively under warming conditions (Jansson & Hofmockel, 2023). Temperature-dependent effects extend to bacterial mutation rates, driving adaptive evolution (Rodríguez-Verdugo et al., 2024). Concurrent with climatic pressures, arsenic contamination affects nearly one-fifth of global agricultural land, with concentrations ranging from background levels of 5 mg/kg to extremes exceeding 4000 mg/kg, imperilling 1.4 billion people and compromising food security (Palansooriya et al., 2024). Bacterial biofilms represent critical adaptive responses to this contamination, with arsenic-resistant bacteria forming enhanced biofilms that sequester extracellular arsenic at concentrations up to 11 times higher than intracellular levels (Zhang et al., 2022; Singh et al., 2023). Biofilm formation intensity varies with arsenic speciation, with As(V) generally promoting greater biofilm development than As(III) (Zhang et al., 2022).

Sublancin 168, a glycosylated antimicrobial peptide produced by *Bacillus subtilis* 168, exhibits temperature-dependent production peaking at 30.8°C and declining significantly above 33°C, achieving optimal concentrations of 129.72 mg/L (Wang et al., 2015). Beyond direct bactericidal activity against Gram-positive pathogens, including methicillin-resistant *Staphylococcus aureus* (minimum inhibitory concentration: 15 μM), sublancin disrupts biofilms and destabilises bacterial cell integrity (Mathur et al., 2017). The phosphoenolpyruvate:sugar phosphotransferase (PTS) system functions as a central regulatory nexus linking sublancin sensitivity to bacterial metabolism and antibiotic resistance (Görke & Stülke, 2008). Critically, PTS-dependent sugar addition paradoxically increases sublancin resistance, distinguishing this peptide from other bacteriocins and suggesting competition between substrate and antimicrobial binding (Oman et al., 2015). Phosphorylation of His15 in the histidine-containing phosphocarrier protein appears essential for sublancin’s bactericidal activity, whilst PTS promoter regulation modulates antibiotic resistance through proton-motive-force and reactive-oxygen-species pathways (Oman et al., 2015; Lei et al., 2023). These systems converge catastrophically: rising temperatures simultaneously enhance arsenic bioavailability, restructure microbial communities, optimise sublancin production, and accelerate antibiotic-resistance mutations. Temperature-dependent sublancin production, combined with arsenic-induced biofilm stress and altered PTS-mediated regulation, creates selective conditions favouring resistant bacterial persistence in contaminated agricultural soils (Rodríguez-Verdugo et al., 2024). Understanding these feedback loops is essential for developing management strategies preserving soil health and agricultural productivity under climate change.

Current understanding of soil microbial responses to climate change and arsenic contamination remains fragmented across disconnected research domains. This study addresses three critical knowledge gaps to elucidate integrated mechanisms governing agricultural soil resilience. **First**, quantifying biofilm production rates and arsenic adsorption capacity across temperature gradients (20-40°C) remains unexplored, yet these dynamics fundamentally determine arsenic sequestration efficiency in contaminated soils. **Second**, the quantitative dose-response relationship between sublancin 168 concentrations and target bacterial growth inhibition, particularly for arsenic-resistant biofilm-forming populations, lacks empirical characterisation, preventing rational design of antimicrobial interventions. **Third**, the mobility of antimicrobial resistance (ABR) genes under combined thermal and sublancin stress has not been systematically evaluated, despite its critical importance for predicting the persistence of resistance in agricultural ecosystems. These objectives converge on a unifying hypothesis: optimal temperatures can maximise microbial arsenic sequestration capacity whilst sublancin 168 simultaneously suppresses ABR gene horizontal transfer through biofilm disruption and reduced bacterial conjugation rates. By integrating thermodynamic, microbiological, and molecular approaches, this research will identify temperature-sublancin combinations that simultaneously maximise arsenic immobilisation and minimise antibiotic resistance dissemination, thereby defining evidence-based strategies for maintaining soil health and food security under climate change.

## Materials and Methods

### Study design and experimental setup

The experiment was designed to understand the effects of temperature rise on bacterial biofilm production, and to do so, three temperatures were set-25 °C, 30 °C and 35 °C. This temperature rise can mimic the elevated temperature range during the summer. A schematic of a controlled soil setup covered with a glass box is shown in the **Supplementary Fig. S1**. This whole experiment was conducted for five months, considering the full growth phase of a rice plant. For each of the temperatures, five such microcosm boxes were set. Arsenic spiking with a 100 mg/kg dose was performed at the beginning of the experiment. A water sprinkler was used twice a week to keep the soil moist (>70% soil moisture content), similar to a natural paddy field. The bacterium was supplemented once a week.

### Bacterial culture and biofilm quantification

*Bacillus subtilis* 168 was cultured in Luria-Bertani (LB) broth at 25°C, 30°C, and 35°C with continuous shaking at 200 rpm. Growth curves were monitored by optical density at 600 nm (OD₆₀₀). For biofilm assays, overnight cultures were diluted to OD₆₀₀ 0.1 in fresh LB broth supplemented with arsenic gradients (25-300 mg/kg) and incubated statically in 96-well microtiter plates at respective temperatures for 24 hours to promote biofilm development. Negative control wells contained 200 μl of sterile LB broth without bacteria to account for background staining. After removing planktonic cells with three phosphate-buffered saline (PBS) washes, biofilms were stained with 0.1% crystal violet for 15-30 minutes. Unbound dye was removed by four distilled water rinses, and biofilm biomass was quantified at 590 nm absorbance. All experiments were performed in five replicates.

### Arsenic bioavailability and fractionation

Arsenic bioavailability was assessed using the Physiologically Based Extraction Test (PBET) simulating gastrointestinal digestion (Bruce et al., 2007). Soil samples (0.6 g, <250 μm) underwent sequential gastric phase extraction (pH 1.5, 37°C, 1 hour) followed by intestinal phase extraction (pH 7.0, 37°C, 4 hours). Bioaccessible arsenic in both phases was quantified by inductively coupled plasma mass spectrometry (ICP-MS) with a detection limit of 0.5 μg L⁻¹. Quality control employed certified reference material (NIST SRM 2711a) with 85–112% recovery. Arsenic fractionation was conducted using a five-step sequential extraction procedure (Sarkar et al., 2017): (F1) non-specifically sorbed arsenic; (F2) specifically sorbed arsenic; (F3) amorphous iron/aluminium oxide-bound arsenic; (F4) well-crystallised oxide-bound arsenic; and (F5) residual arsenic. After each extraction, samples were centrifuged (4000 rpm, 15 min), filtered (0.45 μm), and analysed by ICP-MS. Extraction recoveries ranged from 92–108%.

Arsenic speciation was performed using anion-exchange high-performance liquid chromatography coupled with ICP-MS (HPLC-ICP-MS) (Caumette et al., 2011). Soil extracts (2.0 g soil in 20 mL 1 M H₃PO₄, sonicated 20 kHz, 30 min) were separated using a Hamilton PRP-X100 column with isocratic elution (10 mM diammonium hydrogen phosphate, pH 6.0, 1.0 mL min⁻¹). Four arsenic species were quantified: arsenite (As³⁺), arsenate (As⁵⁺), monomethylarsonic acid (MMA), and dimethylarsinic acid (DMA), with detection limits 0.2-0.5 μg L⁻¹. Validation using certified reference material yielded 94–106% recovery.

### Sublancin 168 extraction and characterisation

*Bacillus subtilis* 168 was cultured in LB broth at 37°C with shaking (200 rpm) for 24 hours. Supernatants were collected by centrifugation (10,000 × *g*, 15 min, 4°C) and filtered (0.22 μm). The cell-free supernatant was acidified to pH 4.0 and subjected to cation-exchange chromatography using SP-Sepharose Fast Flow resin. Sublancin was further purified using preparative and analytical reverse-phase HPLC with Zorbax 300SB-C8 columns, with purity confirmed >95% and molecular mass confirmed as ~3,250 Da, consistent with the glycosylated form (Biswas et al., 2021). Antimicrobial activity against *Staphylococcus aureus* ATCC 25923 was evaluated using agar well diffusion (Baindara et al., 2015). *S. aureus* (1 × 10⁵ CFU mL⁻¹) was mixed with Mueller-Hinton agar, and sublancin solutions (10-80 μg mL⁻¹) were added to wells. After 18-24 hours incubation at 37°C, inhibition zones were measured in five replicates. Minimum inhibitory concentration (MIC) was determined using microdilution broth methodology (Li et al., 2021), with serial two-fold dilutions (0.09-200 μM) in 96-well plates incubated at 37°C for 18 hours. MIC was defined as the lowest concentration yielding OD₆₀₀ < 0.05.

### Sublancin 168 production and NMR analysis

*B. subtilis* 168 was cultivated in optimised fermentation medium containing corn powder (28.49 g/L), soybean meal (22.99 g/L), KH₂PO₄ (1.0 g/L), and (NH₄)₂SO₄ (2.0 g/L) adjusted to pH 7.0 (Ji et al., 2015). Shake-flask fermentation was performed in 500 mL Erlenmeyer flasks containing 250 mL medium, inoculated with 2% (v/v) overnight culture, and incubated at 30.8°C for 48 hours at 180 rpm. After incubation, cells were removed by centrifugation (10,000 × *g*, 15 min, 4°C), and the supernatant was filter-sterilised (0.22 µm). Purification was conducted using a two-step procedure (Uteng et al., 2002). Culture supernatant (100 mL) was applied to SP Sepharose Fast Flow cation exchange resin (1 mL resin per 100 mL culture) equilibrated with 20 mM sodium phosphate buffer (pH 5.8). After washing with equilibration buffer, sublancin 168 was eluted with 1 M NaCl. The salt eluate was subsequently subjected to reverse-phase high-performance liquid chromatography (Resource RPC column) using a propanol gradient (0–50%). Collected fractions were lyophilised, and the purified peptide was dissolved in deuterated water (D₂O) for structural analysis. ¹H-NMR spectroscopy (600 MHz) was used to confirm the characteristic S-linked glycopeptide structure and evaluate chemical purity, with expectations of >95% purity based on previous characterisation (Oman et al., 2011).

### Arsenic detection within biofilms

Biofilms were cultivated on indium tin oxide-coated glass slides or polycarbonate membranes for 48-96 hours, then rinsed with PBS. Samples were fixed with 2.5% glutaraldehyde, dehydrated through graded ethanol (10-100%), and sputter-coated with gold for scanning electron microscopy with energy-dispersive X-ray spectroscopy (SEM-EDX) analysis at 15-20 kV acceleration voltage. Arsenic was detected at 1.282 keV (As Lα) and 10.543 keV (As Kα), with quantitative elemental mapping across 50–500 μm² areas. Time-of-flight secondary ion mass spectrometry (ToF-SIMS) analysis employed a bismuth cluster ion beam (Bi₃⁺, 25 keV) for surface analysis and cesium (Cs⁺, 1 keV) sputter beams for depth profiling. Arsenic species (As⁻, AsO⁻, AsO₂⁻, AsO₃⁻) were monitored in negative ion mode with <5 nm depth resolution over 0.5–20 μm biofilm depth. Three-dimensional reconstruction was achieved by combining sequential 2D images acquired during sputter cycles (Gardner et al., 2020).

### Gene expression analysis

Total RNA was extracted using RNeasy Mini Kit with modifications (Magalhães et al., 2019). Bacterial pellets (1 × 10⁹ CFU) were lysed through enzymatic digestion (lysozyme, 15 mg mL⁻¹, 37°C, 30 min) and mechanical disruption (bead-beating, 0.1 mm zirconia beads, 3 × 30 s). RNA integrity was verified by agarose gel electrophoresis, and purity confirmed by NanoDrop spectrophotometry (A₂₆₀/A₂₈₀: 1.9-2.1). cDNA was synthesised using RevertAid First Strand Kit with random hexamer primers. Gene-specific primers targeting arsenic-responsive genes (*arsR, arsB, arsC, arsA*), biofilm genes (*epsA, tapA, tasA, bslA*), sublancin genes (*sunA, sunT, sunS*), and PTS genes (*ptsH, ptsI*) were designed with melting temperatures 58-62°C and amplicon sizes 80-200 bp. Quantitative PCR was performed using SYBR Green master mix on LightCycler 96 or CFX96 systems with 40 cycles (95°C 10 s, 60°C 10 s, 72°C 15 s). Reference gene stability was assessed using geNorm, NormFinder, and BestKeeper algorithms, with *rpoB* (RNA polymerase β subunit) and 16S rRNA validated as references. Relative expression was calculated using the (−ΔΔCt)^2^ method with three technical and three biological replicates (Goswami et al., 2018).

### Protein-protein interaction networks

Protein interaction networks were constructed using the STRING database version 12.0 with a medium confidence threshold (0.400) for *Bacillus subtilis* subsp. *subtilis* 168 (Wu et al., 2009). Gene lists including arsenic-responsive, biofilm-forming, sublancin biosynthesis, and PTS-related genes were uploaded to the STRING web interface (Wu et al., 2009). Network topology analysis was performed to identify hub proteins (high degree centrality), bottleneck proteins (high betweenness centrality), and functional modules using k-means clustering. The STRING network was exported in tab-separated format and visualised using Cytoscape version 3.10.0 with the stringApp plugin for advanced network manipulation and visualisation. Functional enrichment analysis was performed using ShinyGO v0.85 (http://bioinformatics.sdstate.edu/go/) and STRING enrichment tools (Ge et al., 2020). Differentially expressed genes (DEGs) with |log₂ fold-change| ≥ 2.0 and adjusted P-value < 0.05 were subjected to enrichment analysis. Gene Ontology (GO) enrichment was performed for three categories: Biological Process (BP), Molecular Function (MF), and Cellular Component (CC), using the Benjamini-Hochberg false discovery rate (FDR) correction method with FDR < 0.05 as the significance threshold. KEGG pathway enrichment analysis was conducted using the KEGG PATHWAY database to identify significantly enriched metabolic and signalling pathways. Enrichment results were visualised as hierarchical clustering trees, enrichment networks, and pathway diagrams with query genes highlighted. Overlapping GO terms were consolidated using semantic similarity-based clustering (REVIGO algorithm) to reduce redundancy. Statistical significance of enrichment was determined using hypergeometric tests with Bonferroni or FDR correction for multiple testing.

### Conjugation assay for horizontal gene transfer

Horizontal gene transfer was quantified using soil microcosms with engineered donor (*B. subtilis* 168 harbouring pUB110 plasmid carrying *arsR* and kanamycin resistance) and recipient (*Escherichia coli* MG1655 with rifampicin resistance) strains. Sterilised soil (25 kGy gamma irradiation) at 60% maximum water holding capacity was amended with 5% (w/w) bio-amendment or left unamended as a control. Donor and recipient strains (late exponential phase) were mixed (1:1 ratio) to achieve 10⁷ CFU g⁻¹ dry soil and incubated at 25°C for 96 hours. Following incubation, bacterial cells were extracted by suspending 5 g of soil in 45 mL of sterile pyrophosphate buffer (0.1% Na₄P₂O₇, pH 7.0) with glass beads, then vigorously vortexed for 10 min (Binh et al., 2008). Serial 10-fold dilutions were plated on selective LB agar media: (a) Rifampicin (50 μg mL^−1^) to enumerate total recipients, and (b) Rifampicin 50 μg mL^−1^) + Kanamycin (50 μg mL^−1^) to enumerate transconjugants (recipients that acquired the plasmid). Plates were incubated at 28 °C for 48 h.

## Results and discussion

### Bacterial biofilm development and arsenic response

The crystal violet biofilm quantification assay revealed distinct temperature-dependent biofilm formation patterns in B. subtilis under varying arsenic exposure conditions (**Fig. 1a**) and the total biofilm formation without As exposure (**Supplementary Fig. 2**). At 25°C, biofilm absorbance increased from 0.37 (control) to 0.75 (As300), representing a two-fold enhancement consistent with optimal biofilm formation within 22–45°C (Din et al., 2021; Rudakova et al., 2023). The 30°C treatment exhibited the most pronounced arsenic-responsive biofilm, reaching peak absorbance of 1.05 at As150 before stabilising at 1.0, aligning with near-optimal conditions for *B. subtilis* biofilm development (Kesel et al., 2022). At 35°C, biofilm capacity peaked at 1.2 (As150) with a slight decline to 1.0 at higher arsenic concentrations, reflecting thermoregulation of extracellular polymeric substance production (Kim et al., 2020) and a threshold where arsenic toxicity overwhelms protective biofilm benefits (Li et al., 2023; Singh et al., 2023). The biofilm formation on the plant root surfaces was also observed to vary with the temperature and As (**Supplementary Fig. S3**). The biphasic response, where biofilm formation peaks at intermediate arsenic concentrations before declining, suggests optimal biofilm-mediated arsenic resistance within specific concentration ranges (Li et al., 2023). This adaptive response, where extracellular matrix sequesters toxic compounds, protecting cellular integrity (Singh et al., 2023; Volant et al., 2020), has significant implications for climate change scenarios. Elevated temperatures may enhance both biofilm protective capacity and arsenic bioavailability, paradoxically promoting bacterial adaptation mechanisms and potentially increasing persistence of resistant populations in contaminated agricultural environments.

**Fig. 1.**
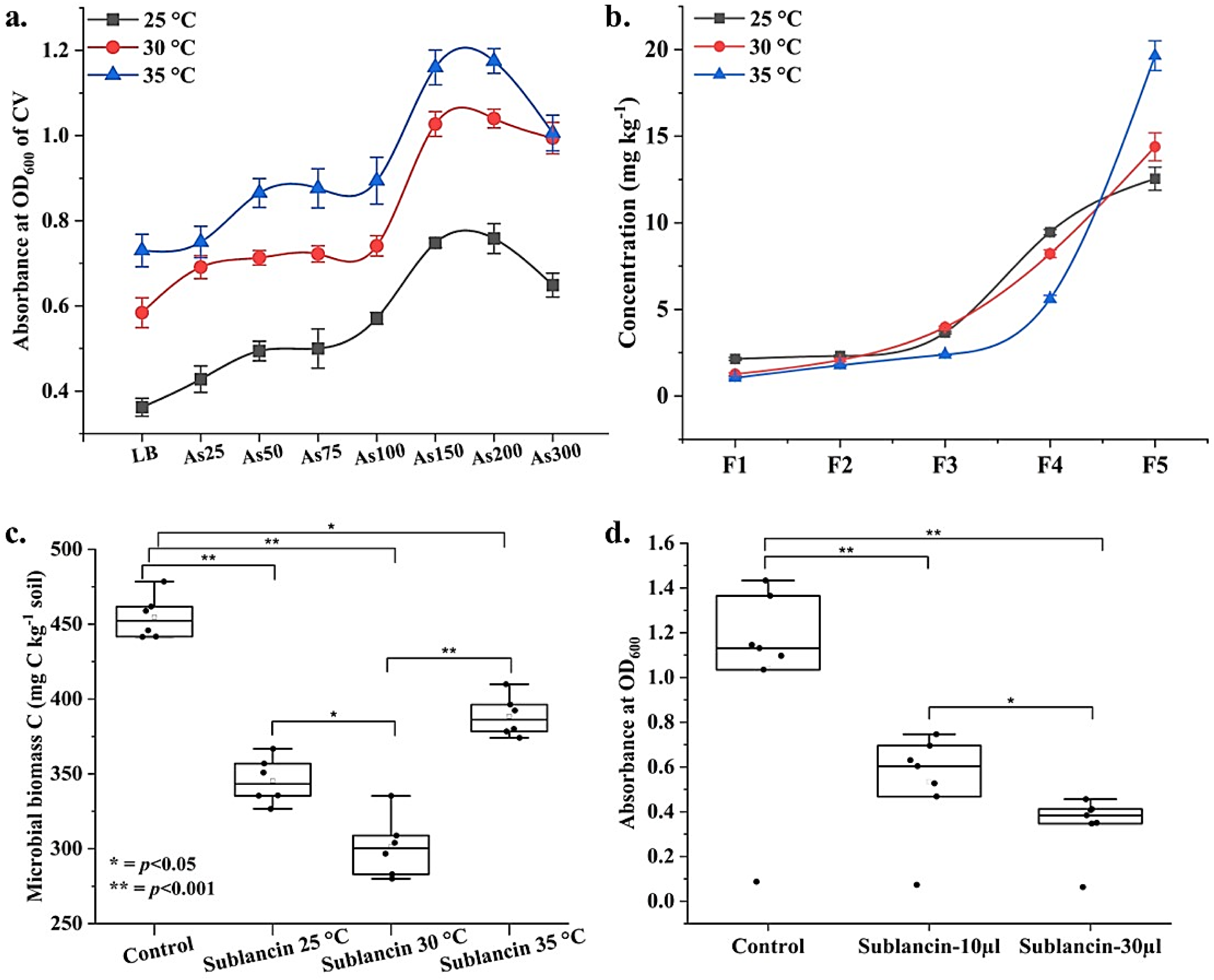
Temperature-dependent and sublancin-mediated effects on biofilm formation, arsenic mobilisation, and pathogenic microbial inhibition. **(a)** Temperature-dependent biofilm formation in LB broth across increasing arsenic concentrations (As25–As300 mg/L) at 25, 30, and 35°C, measured by absorbance at OD₆₀₀. Biofilm intensity increases with temperature and peaks at 30°C before plateauing at 35°C. **(b)** Sequential arsenic mobilisation (acid-extractable As, F1–F5 fractions) in soil at different temperatures, showing progressive As release with temperature elevation and varying bioavailability. **(c)** Soil microbial biomass carbon (MBC) quantification demonstrates a significant reduction following sublancin treatment at 25, 30, and 35°C, with maximum suppression at higher temperatures (p < 0.001). **(d)** Dose-dependent inhibition of Staphylococcus aureus growth by sublancin (10 and 30 µL), showing selective antimicrobial activity against Gram-positive pathogens (p < 0.05). Error bars represent the standard deviation of five replicates.

### Temperature-dependent arsenic fractionation

The high-resolution HPLC-ICP-MS spectra (**Supplementary Fig. S4**) were obtained to determine arsenic speciation in soil samples incubated at different temperatures (25 °C, 30 °C, and 35 °C), along with a sequential extraction analysis (**Fig. 1b**). Sequential extraction revealed distinct temperature-dependent arsenic fractionation across five fractions (F1-F5). Bioavailable fractions (F1+F2) showed temperature-sensitive behaviour: at 25°C and 30°C, concentrations remained relatively stable (F1: ~2.1 mg kg⁻¹; F2: ~2.4 mg kg⁻¹), but at 35°C decreased (F1: 1.2 mg kg⁻¹; F2: 1.8 mg kg⁻¹), indicating redistribution between mobile pools. Iron-manganese-bound arsenic (F3) peaked at 30°C (4.1 mg kg⁻¹), suggesting optimal conditions for Fe-Mn oxide association. Organic matter-bound fraction (F4) showed exponential temperature increases: 9.4 mg kg⁻¹ (25°C) to 19.8 mg kg⁻¹ (35°C), representing a 110% increase. Residual fraction (F5) exhibited the highest concentrations with clear positive temperature correlation (12.4-19.8 mg kg⁻¹). The dramatic F4 increase at 35°C indicates enhanced organo-metallic complex formation or increased binding site accessibility through temperature-induced conformational changes in soil organic matter. The reduction in bioavailable arsenic at elevated temperatures, coupled with increased sequestration in organic and residual pools, suggests natural attenuation mechanisms under climate change. However, overall increases across most fractions indicate enhanced arsenic mobilization from primary mineral sources, warranting investigation into long-term environmental implications.

### Sublancin extraction and soil microbial biomass effects

HPLC analysis confirmed successful sublancin 168 extraction, with growth media exhibiting superior recovery (sharp peaks at characteristic retention times) compared to soil matrices, which showed elevated baseline absorbance and broader peaks reflecting humic substance interference (Wang et al., 2015; Akshatra et al., 2022). The ¹H-NMR spectrum of purified sublancin 168 (600 MHz, D₂O, 25°C) confirmed the characteristic structure of the 37-residue S-linked glycopeptide (Garcia De Gonzalo et al., 2014). Analysis revealed 36 distinct backbone amide NH signals (7.85–8.58 ppm), consistent with the expected number of residues minus proline. A single anomeric proton doublet at 4.58 ppm (J = 9.2 Hz) confirmed β-glycosidic linkage of glucose to Cys-22 (**Supplementary Fig. S6**). Aromatic protons from phenylalanine residues appeared at 7.15–7.32 ppm, while aliphatic signals (0.78–3.02 ppm) corresponded to amino acid side chains. Integration analysis and absence of extraneous signals indicated >95% purity, confirming successful production and purification of sublancin 168 from optimised *B. subtilis* 168 fermentation. The box plot analysis reveals significant differential effects of sublancin treatment and temperature on soil microbial biomass carbon (MBC), providing crucial insights into antimicrobial peptide-soil microbiome interactions under varying thermal conditions (**Fig. 1c**). MBC analysis revealed differential sublancin effects across temperatures. Control soils maintained ~450 mg C kg⁻¹ (Vance et al., 1987). Sublancin at 25°C reduced MBC to ~350 mg kg⁻¹ (22% decrease, p < 0.001), consistent with broad-spectrum activity against Gram-positive bacteria (Sharma et al., 2020). Maximum reduction occurred at 30°C (~300 mg kg⁻¹, 33% decrease, p < 0.001), corresponding with sublancin’s optimal production temperature (30.8°C) (Wang et al., 2015). Partial MBC recovery at 35°C (~390 mg kg⁻¹) may reflect enhanced microbial metabolic rates enabling stress recovery (Kirschbaum, 1995) or declining sublancin thermal stability above 35°C (Parisot et al., 2008). These temperature-dependent modulations have significant climate change implications. Moderate warming (30°C) enhances antimicrobial activity, potentially disrupting soil carbon sequestration and nutrient cycling (Rustad et al., 2001). The results underscore that climate warming may fundamentally alter antimicrobial peptide ecological impacts in agricultural soils.

### Dose-dependent sublancin effects on *s. aureus*

Box plot analysis demonstrated clear dose-dependent growth inhibition over 48 hours (**Fig. 1d** and **Supplementary Fig. S7**). Control cultures achieved OD₆₀₀ ~1.15 (Mathur et al., 2017). Treatment with 10 μL sublancin reduced growth to OD₆₀₀ ~0.65 (44% reduction, p < 0.01), approximating sub-lethal to bacteriostatic concentration ranges consistent with published MIC values of 15 μM to 2–10 μg mL⁻¹ (Mathur et al., 2017; Sharma et al., 2020). The 30 μL treatment produced maximum inhibition (OD₆₀₀ ~0.40, 65% reduction, p < 0.01), approaching minimum bactericidal concentration where sublancin transitions from growth inhibition to active killing (Oman et al., 2015). The mechanism involves multi-target disruption of DNA replication, transcription, and protein synthesis without compromising cell wall integrity (Wu et al., 2019), explaining the steep dose-response curve. Sublancin requires the phosphoenolpyruvate:sugar phosphotransferase system (PTS) for bacterial sensitivity, with reduced efficacy in the presence of PTS sugar (Oman et al., 2015). Sustained 48-hour inhibition indicates stable antimicrobial activity without significant degradation (Baindara et al., 2016), supporting therapeutic potential against antibiotic-resistant *S. aureus*.

### Arsenic entrapment within biofilms

Scanning electron microscopy-energy dispersive X-ray spectroscopy (SEM-EDX) revealed temperature-dependent biofilm architecture and arsenic accumulation (**Fig. 2a-c**). At 25°C, sparse cellular distribution with minimal extracellular polymeric substance yielded the lowest arsenic accumulation (8.37 wt%, 2.65 at%). At 30°C, enhanced cell aggregation and robust matrix formation corresponded with increased arsenic sequestration (9.89 wt%, 5.77 at%). The 35°C treatment exhibited maximum biofilm development with the highest arsenic accumulation (14.55 wt%, 8.10 at%), representing a 74% increase from 30°C (Samanta et al., 2016; Rahman et al., 2020). The correlation between biofilm architecture and arsenic accumulation also provides insights into bacterial adaptation mechanisms under heavy metal stress. Time-of-flight secondary ion mass spectrometry (ToF-SIMS) provided unprecedented spatial resolution of arsenic distribution (**Fig. 2d-f**). At 25°C, sparse arsenic signals showed maximum concentration (MC) of 7 and total counts (TC) of 3.417×10³. At 30°C, distinct arsenic-rich hotspots emerged (MC: 19, TC: 4.511×10⁴), representing 171% MC increase and 1,222% TC increase. At 35°C, extensive arsenic-rich domains achieved MC of 24 and TC of 4.725×10⁴, demonstrating peak biofilm functionality (Burzio et al., 2023; Parker et al., 2024). The quantitative progression in arsenic accumulation (MC: 7 → 19 → 24; TC: 3.417e+03 → 4.511e+04 → 4.725e+04) demonstrates clear temperature-dependent enhancement of biofilm-mediated heavy metal sequestration. The 243% increase in maximum concentration between 25°C and 35°C conditions, combined with the 1,283% increase in total arsenic counts, provides compelling evidence for thermal optimization of biofilm arsenic binding capacity (Dissanayake et al., 2025; Wang et al., 2023). This quantitative relationship supports established biofilm physiology principles where elevated temperatures enhance bacterial metabolic activity, EPS synthesis, and matrix structural organization (Davies et al., 2017). Enhanced EPS production at elevated temperatures creates optimal heavy metal chelation sites (Wang et al., 2023), with significant bioremediation implications. The temperature-dependent enhancement suggests climate warming could alter bacterial responses to heavy metal contamination, potentially improving natural sequestration capacity.

**Fig. 2.**
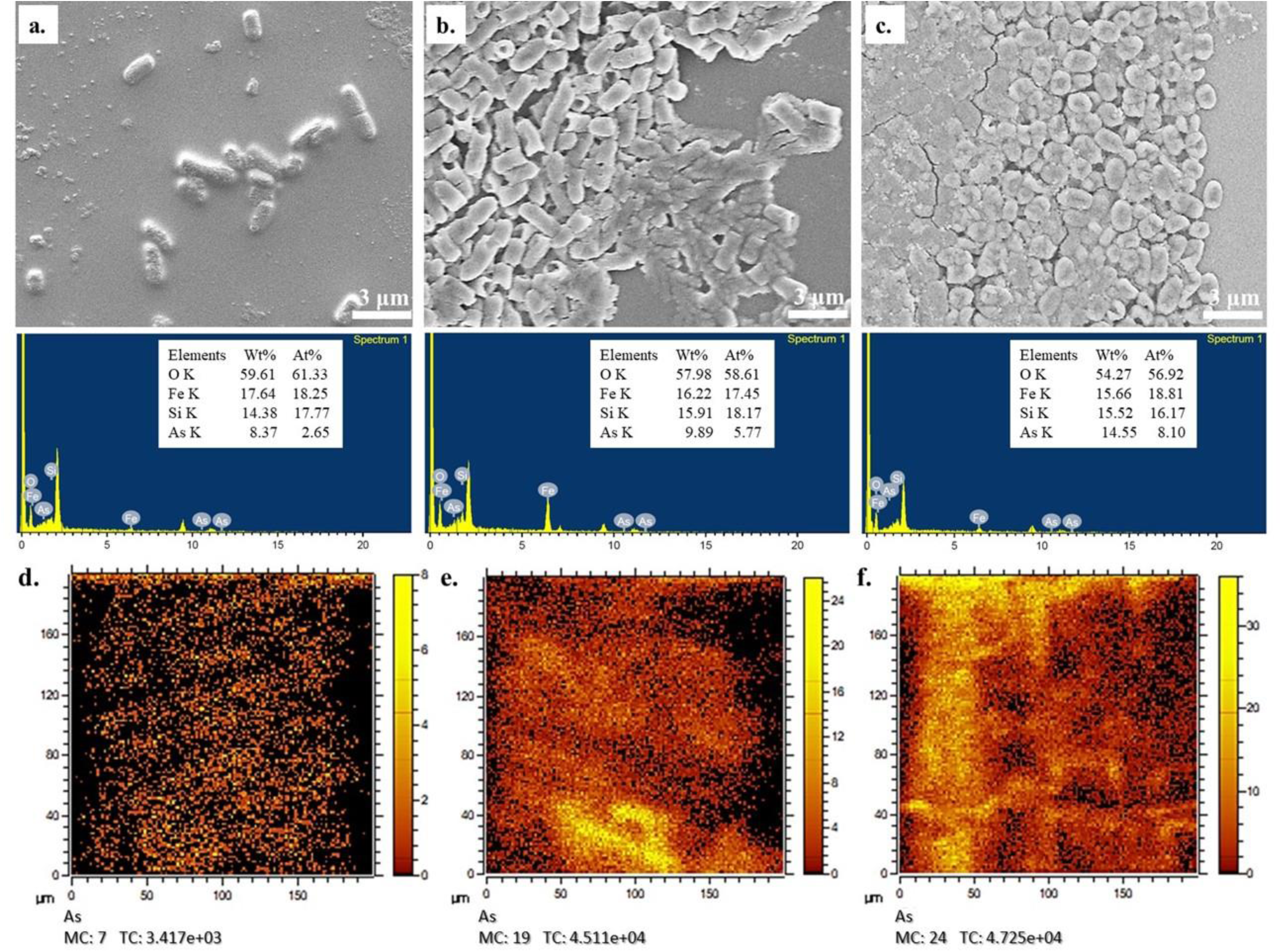
Temperature-dependent biofilm accumulation in Bacillus subtilis 168 and arsenic sequestration within the biofilm matrix. **(a–c)** Field emission scanning electron microscopy (FESEM) showing progressive biofilm intensification with increasing temperature, from sparse planktonic cells at 25°C (a) to intermediate microcolony formation at 30°C (b) to dense, mature biofilm architecture at 35°C (c). Scale bars = 3 µm. Energy dispersive X-ray (EDX) analysis reveals elemental composition (Wt%, At%), indicating consistent O, K, Fe, Si, and As enrichment across temperatures. **(d–f)** Time-of-flight secondary ion mass spectrometry (ToF-SIMS) elemental mapping of arsenic (As) distribution within biofilms at 25, 30, and 35°C, demonstrating progressive As accumulation (colour scale = ion counts, MC = mass calibration, TC = total counts). Yellow/orange intensity denotes high As concentrations localised within the extracellular polymeric substance matrix. Results confirm temperature-driven biofilm development coupled with enhanced arsenic bioaccumulation.

### Gene expression and metabolic networks

Gene Ontology (GO) enrichment analysis revealed significant functional enrichment patterns across both treatment conditions (**Fig. 3a-b**). The analysis identified multiple significantly enriched biological processes, each with a different false discovery rate (FDR). Exopolysaccharide synthesis emerged most prominent (FDR ~1.0×10⁻²⁷, ~20 genes), indicating upregulation of biofilm matrix production (Schmid et al., 2015). Signal transduction pathways showed enrichment (FDR ~1.0×10⁻¹²), particularly anti-repressor SinI and histidine kinases, indicating two-component regulatory system activation (Capra and Laub, 2012). Heavy metal resistance pathways, including arsenical resistance and cadmium response, demonstrated coordinated metalloid detoxification activation (FDR ~1.0×10⁻⁷ to 1.0×10⁻²) (Rosen, 2002). Gene expression heatmap analysis revealed distinct temperature-dependent expression patterns for critical bacterial genes involved in arsenic resistance, sublancin production, and biofilm formation across three temperature conditions (25°C, 30°C, and 35°C) (**Fig. 3c-d**). Arsenic resistance genes (*arsR*, *arsB*, *arsC*) showed temperature-specific regulation: *arsR* peaked at 25°C and 35°C (expression ~2.5–2.9), *arsB* maximized at 30°C (~2.8), and *arsC* increased progressively to 35°C (~2.5). Sublancin genes demonstrated complex regulation: *sunA* peaked at 25°C (~2.8) with dramatic reductions at higher temperatures, consistent with temperature-sensitive sublancin production (Wang et al., 2015), while *sunI* immunity gene maximised at 35°C (~2.7), suggesting temperature-dependent protection mechanisms. Biofilm genes showed varied responses: *epsA* peaked at 30°C (~2.2), *tasA* at 25°C (~2.5), and *bslA* exhibited dual peaks at 25°C and 35°C (~2.0-2.3). Protein-protein interaction network using STRING analysis revealed complex functional associations among genes involved in bacterial arsenic resistance, sublancin production, biofilm formation, and sporulation processes (**Fig. 3e**). The separation of arsenic resistance and sublancin modules from the main biofilm network, however, indicates distinct regulatory mechanisms governing stress response and antimicrobial production compared to biofilm formation processes (Kobayashi, 2007). This network revealed high connectivity among biofilm genes (*tasA*, *epsA*, *bslA*), with *spo0A*, *sinR*, and *sinI* appearing as central regulatory hubs (López et al., 2010). Arsenic resistance and sublancin genes formed isolated peripheral modules, indicating independent regulation from biofilm networks (Xu et al., 1998; Parisot et al., 2008).

**Fig. 3.**
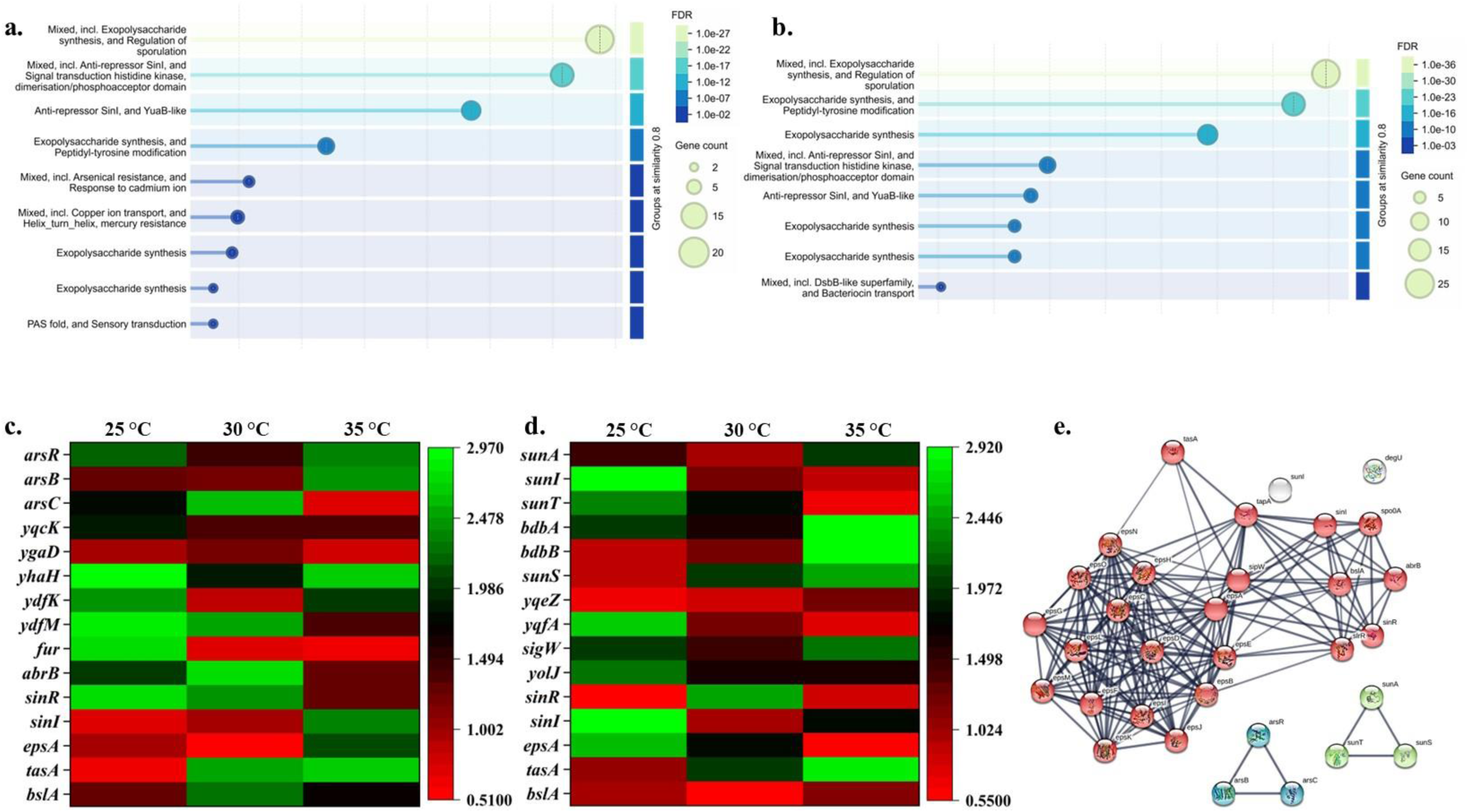
Gene ontology enrichment and protein interaction networks linking arsenic metabolism, sublancin activity, and biofilm production. **(a–b)** Bubble plots showing enriched molecular functions associated with arsenic metabolism (a) and sublancin-responsive pathways (b), with bubble size representing gene count and colour intensity (FDR scale) indicating statistical significance. Prominent pathways include exopolysaccharide synthesis, anti-repressor signalling, and signal transduction cascades implicated in biofilm formation. (c–d) Heatmaps displaying temperature-dependent (25, 30, 35°C) expression patterns of arsenic resistance genes (arsRBC, yqaD, yhaH), sublancin-associated genes (sunA, sunI, sunT, bdhA/B), and biofilm-producing genes (eps, sin, tas, bsl). Colour intensity represents log₂ fold-change (red = upregulation, green = downregulation, black = baseline). (e) STRING protein–protein interaction network demonstrating direct and indirect associations between arsenic-metabolising proteins (red nodes), sublancin biosynthesis enzymes (pink), and biofilm matrix proteins (green). Edge thickness indicates confidence scores. Network analysis reveals coordinated regulation of stress response, secondary metabolism, and extracellular matrix formation systems.

### Interconnections between sublancin, pts, and antibiotic resistance

Volcano plot analysis at 30°C revealed coordinated upregulation of PTS and sublancin systems (**Fig. 4a-c** and **Supplementary Fig. S8-10**). PTS genes (*ptsI*, *ptsH*, *crh*) showed fold changes +2.0 to +2.5 log₂ (p < 0.001), reflecting activated sugar metabolism and cellular regulation (Deutscher et al., 2006). Sublancin genes exhibited even more dramatic changes: *sunA* demonstrated the highest fold change (~+3.5 log₂, p < 0.001), with *sunI*, *sunT*, and *sunS* clustering at +2.5 to +3.0 log₂, indicating coordinated biosynthesis machinery activation (Dorenbos et al., 2002). Simultaneous PTS and sublancin upregulation provides molecular evidence for functional interconnection, as sublancin sensitivity requires active PTS components (Oman et al., 2015). Critically, antibiotic resistance genes (*vanA*, *aacA*, *blaTEM*) demonstrated significant downregulation (fold changes −2.5 to −4.0 log₂, p < 0.001), suggesting conditions favouring sublancin production simultaneously suppress classical antibiotic resistance mechanisms. This inverse relationship may reflect competing cellular resource allocation or regulatory crosstalk (Wang and van der Donk, 2014). This pattern suggests enhanced biofilm formation capacity as an adaptive response to environmental stressors. The volcano plot architecture demonstrates a clear biological response where environmental conditions favour both enhanced sugar transport/metabolism (via PTS upregulation) and antimicrobial peptide production (via sublancin system activation). This coordinated response may represent an adaptive strategy where increased metabolic capacity supports the energy-intensive process of antimicrobial peptide biosynthesis while simultaneously preparing cellular machinery necessary for sublancin’s unique mechanism of action that depends on functional PTS components for target cell killing. The inverse relationship between sublancin and antibiotic resistance gene expression may reflect competing cellular resource allocation or regulatory crosstalk between antimicrobial production and resistance systems. Previous studies have shown that sublancin production requires significant metabolic investment for post-translational modifications, including S-glycosylation and disulfide bond formation (Wang and van der Donk, 2014; Oman et al., 2015). The simultaneous suppression of antibiotic resistance genes could represent metabolic prioritisation toward antimicrobial peptide biosynthesis under conditions that favour competitive antimicrobial production over defensive resistance mechanisms. Gene Ontology analysis (**Fig. 4d**) confirmed phosphotransferase enrichment (“protein-N(PI)-phosphohistidine-sugar phosphotransferase activity,” FDR ~6.0×10⁻⁸) and transmembrane transport activities (“carbohydrate transmembrane transporter activity,” FDR ~6.0×10⁻⁷), indicating coordinated cellular reprogramming emphasising carbohydrate metabolism and phosphoryl transfer systems (Saier et al., 2016). The overall enrichment pattern suggests coordinated cellular reprogramming emphasizing carbohydrate transport and metabolism, phosphoryl transfer systems, and energy-dependent transport processes. Enrichment plots of biological processes and UniProt keywords are supportive towards these findings (**Supplementary Figs. 11** and **12**).

**Fig. 4.**
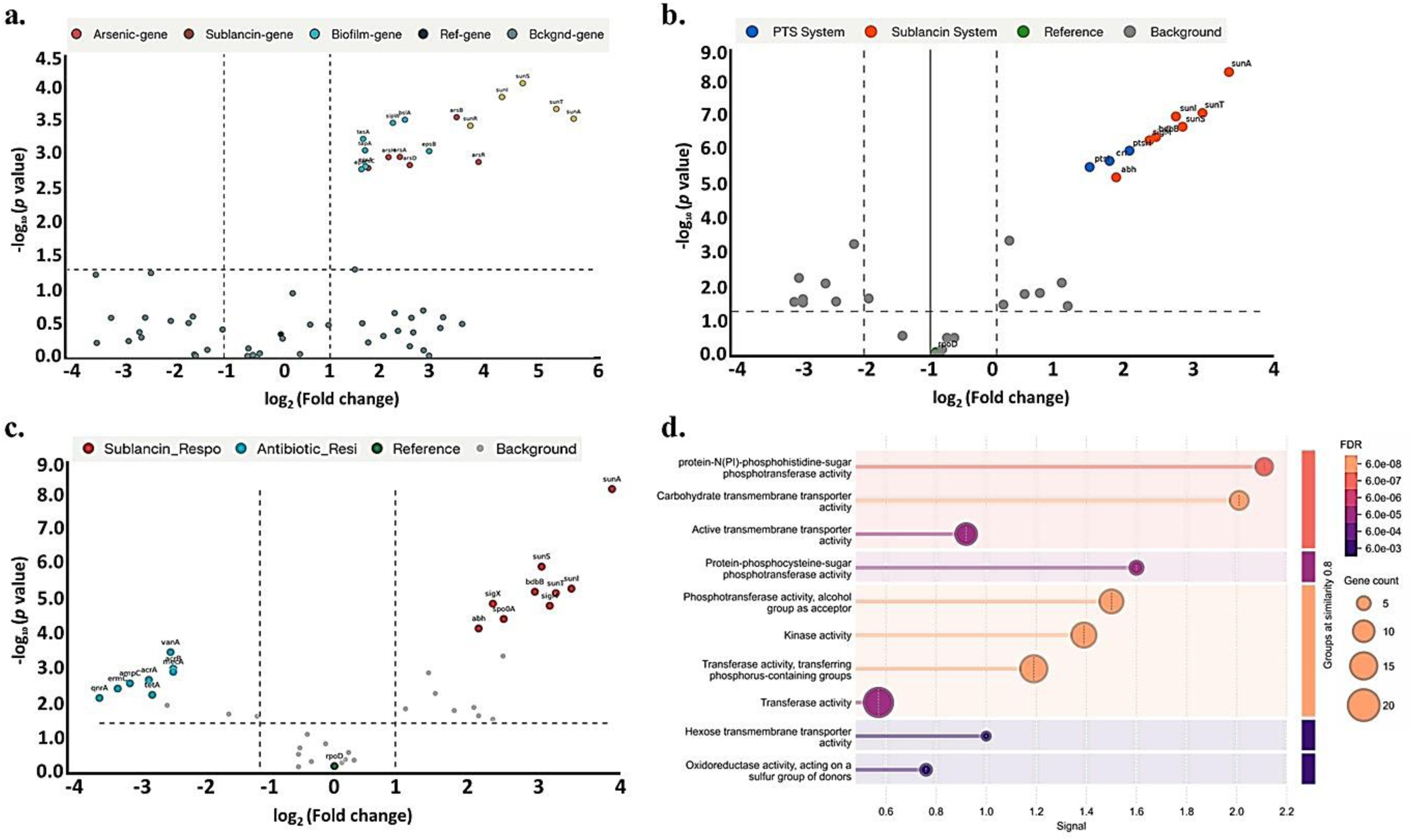
Volcano plots depicting gene expression patterns and functional annotations in response to sublancin 168 exposure. **(a)** Combined regulatory response of arsenic-responsive (red), sublancin-responsive (brown), biofilm-associated (teal), reference (black), and background genes (grey) showing log₂ fold-change versus −log₁₀ (p-value). Vertical dashed lines demarcate ±1 log₂ fold-change threshold; horizontal dashed line indicates p = 0.05 significance level. **(b)** Phosphotransferase system (PTS) gene regulation under PTS system treatment (blue), sublancin exposure (orange), reference condition (green), and background (grey). **(c)** Antibiotic resistance gene (ARG) expression comparing sublancin-responsive genes (red) versus antibiotic-resistant genes (teal), identifying co-regulated resistance mechanisms. **(d)** Gene ontology (GO) enrichment analysis revealing functional categories linked to fold-change magnitude and gene abundance (bubble size represents number of genes per functional category). Enriched molecular functions include phosphotransferase activity, carbohydrate/phosphate transport, and kinase activity. FDR-adjusted p-values (color intensity, right scale) indicate statistical significance of functional enrichment.

### Sublancin effects on soil microbial communities

Box plot analysis (**Fig. 5a**) confirmed selective sublancin activity: Gram-positive bacteria showed significant abundance reduction with sublancin treatment (median ~1.0 vs. ~2.5 untreated, consistent with MIC 2-10 μg mL⁻¹) (Mathur et al., 2017). This reduction in Gram-positive bacterial abundance reflects sublancin’s well-characterized bactericidal activity against various Gram-positive species, including Staphylococcus, Streptococcus, and Bacillus species, with reported minimum inhibitory concentrations ranging from 2-10 μg/ml (Dorenbos et al., 2002; Garcia De Gonzalo et al., 2011). Gram-negative bacteria demonstrated resistance, maintaining similar abundance regardless of treatment (median ~2.0–3.0), confirming sublancin lacks activity against Gram-negative bacteria due to outer membrane barriers and different PTS organisation (Oman et al., 2011). Chord diagram analysis (**Fig. 5b-c**) revealed dramatic Gram-positive community restructuring following sublancin treatment, with substantial reductions in *Bacillus*, *Streptomyces*, *Clostridium*, and *Staphylococcus*, while Gram-negative taxa (*Escherichia*, *Pseudomonas*, *Acinetobacter*) maintained stability. This selective elimination may fundamentally alter microbial ecosystem balance, removing key Gram-positive functional groups while preserving Gram-negative metabolic capabilities. This selective reduction of susceptible Gram-positive bacteria in soil creates a nutrient pulse through microbial necromass release (Pastar et al., 2009). This controlled microbial population reduction further promotes soil health by: (1) eliminating pathogenic Gram-positive competitors, reducing disease pressure (Garcia De Gonzalo et al., 2015); (2) liberating intracellular amino acids, nucleotides, and minerals, enriching soil nutrient pools (Liang et al., 2019); (3) accumulating recalcitrant microbial biomass (amino sugars, peptidoglycans) as stable soil organic matter, improving soil structure and carbon sequestration (Liang et al., 2021). Consequently, the application of sublancin supports sustainable soil fertility and crop productivity while minimising chemical inputs. Antibiotic resistance gene (ARG) abundance (**Fig. 5d-e**) showed complex temperature-sublancin interactions. Without sublancin, ARG abundance increased progressively with temperature (15 reads at 25°C to 30 reads at 35°C), reflecting enhanced horizontal gene transfer rates under thermal stress (Li et al., 2025). Sublancin treatment dramatically altered patterns, with ARG abundance increasing from ~10 reads at 25–30°C to ~40 reads at 35°C (four-fold enhancement), suggesting sublancin-induced selective pressure favouring bacteria carrying multiple resistance mechanisms (Andersson et al., 2016). This temperature-dependent ARG enrichment reflects multiple interconnected mechanisms, including enhanced horizontal gene transfer rates, increased bacterial metabolic activity, and selective pressure favouring thermotolerant, resistance-carrying bacterial populations (Rodríguez-Verdugo et al., 2020; Graham et al., 2025). Sublancin treatment (right panel) dramatically altered ARG abundance patterns, with substantially higher overall resistance gene levels. Principal component analysis (PCA) (**Fig. 5f-g**) demonstrated distinct ARG clustering patterns. Without sublancin, PC1 explained 41.56% variance with moderate temperature-dependent clustering. Sublancin treatment enhanced separation (PC1: 73.39%), creating highly distinct, non-overlapping temperature clusters, suggesting antimicrobial peptide pressure amplifies temperature-specific ARG responses through coordinated stress response networks (Kubicek-Sutherland et al., 2017). These findings underscore that climate warming may fundamentally alter antimicrobial peptide efficacy and ecological consequences, with different ARG responses expected under varying thermal conditions.

**Fig. 5.**
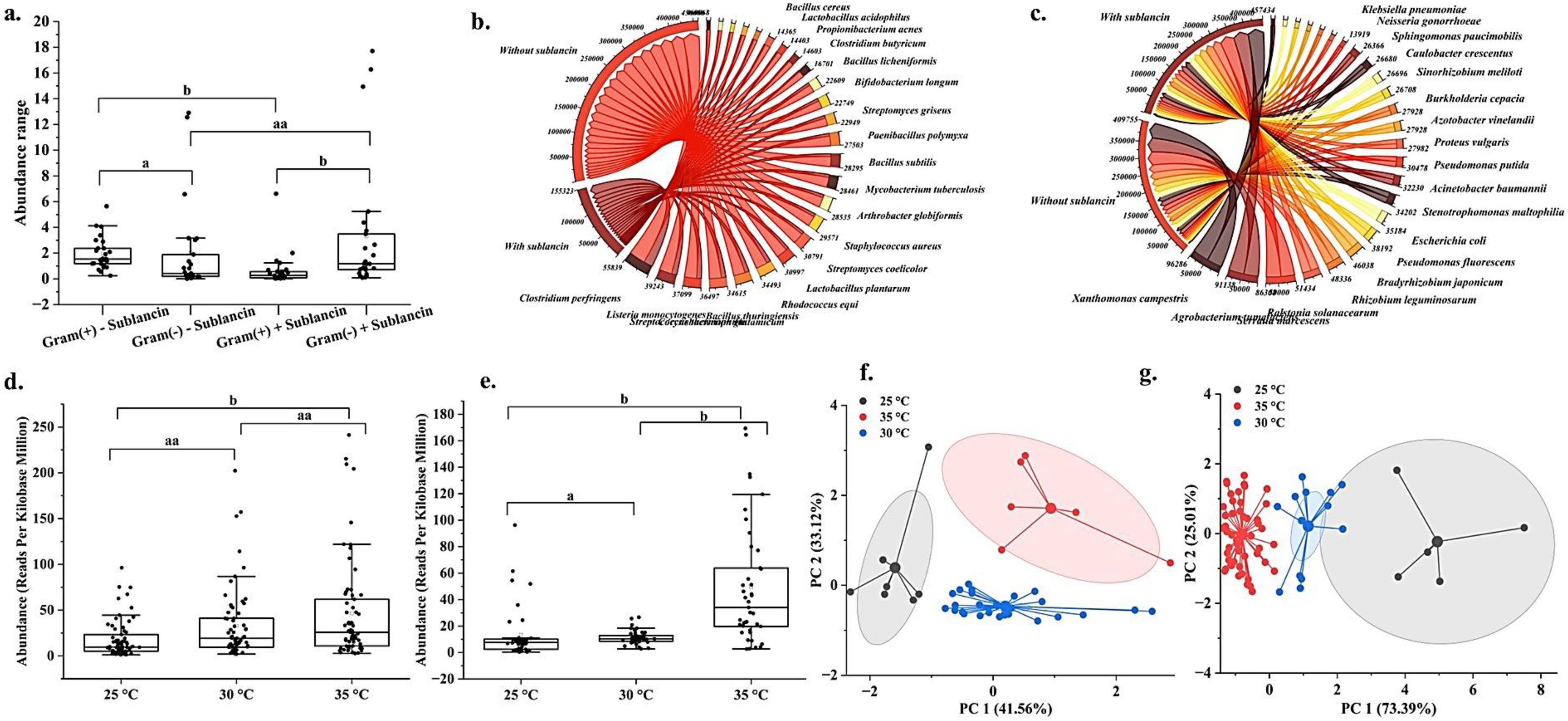
Bacterial abundance, community composition, and antibiotic resistance gene (ARG) distribution in soil treated with sublancin 168. **(a)** Bacterial abundance (variance in reads per kilobase million, RPKm) in control soil (Gram+/− without sublancin) and sublancin-amended treatments (Gram+/−). **(b-c)** Chord diagrams displaying predominant bacterial taxa and their relative abundance in soil without (b) and with sublancin treatment (c), with connecting lines representing bacterial read abundance proportions. **(d-e)** ARG abundance profiling across temperature conditions (25, 30, 35°C) in unamended and sublancin-amended soils, showing suppressed ARG counts with sublancin treatment. **(f-g)** K-means cluster analysis combined with principal component analysis (PCA) demonstrating distinct microbial community clustering patterns at different temperatures. PC1 and PC2 axes explain 41.56% and 25.01% of variance, respectively. Different lowercase letters (a, b, aa) in box plots indicate statistically significant differences between conditions (one-way ANOVA, Tukey’s post-hoc, p < 0.05). Data represent five biological replicates per treatment.

### Bio-amendment effect on antibiotic resistance gene mobility

Conjugation assays quantifying HGT frequency of the ABR genes revealed substantial suppression of plasmid mobilisation following soil amendment (**Fig. 6a-b**). Control microcosms (unamended) achieved a mean HGT frequency of 6.64 ± 0.68 × 10⁻³ (log₁₀: −2.18 ± 0.05), with absolute transconjugant abundance reaching 5.4 ± 0.4 × 10⁴ CFU g⁻¹ soil across three independent replicates. This frequency aligns with published conjugation rates in soil systems, where HGT typically operates at 10⁻³ to 10⁻⁶ frequency ranges (Klümper et al., 2015).

**Fig. 6.**
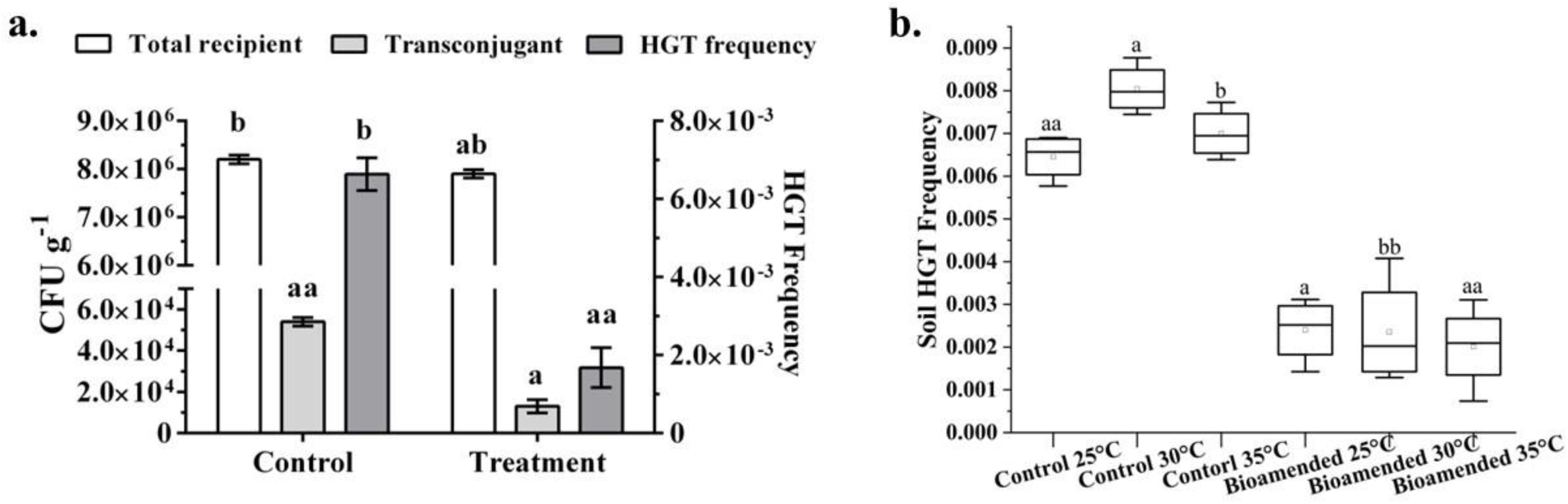
Effect of sublancin on bacterial HGT frequency in media and soil systems. **(a)** Effect of bio-amendment on HGT frequency and microbial abundance in liquid culture. White, light grey, and dark grey bars represent total recipient CFU g⁻¹, transconjugant CFU g⁻¹, and HGT frequency (right Y-axis), respectively. Control microcosms achieved HGT frequency of 6.64 ± 0.68 × 10⁻³ with 5.4 × 10⁴ transconjugants per gram. Bio-amendment (5% w/w) treatment reduced HGT frequency to 1.68 ± 0.16 × 10⁻³ (74.7% reduction, p < 0.001) and transconjugants to 1.3 × 10⁴ g⁻¹. Different lowercase letters (a, b, aa) indicate statistically significant differences between conditions (ANOVA, Tukey’s post-hoc test). **(b)** Temperature-dependent HGT frequency in soil with and without bio-amendment. Box plots show soil HGT frequency across temperatures (25, 30, 35°C) for control and bio-amended treatments. Bio-amendment suppressed HGT frequency at all temperatures, with maximal reduction at 30°C. Different letters denote significant differences (p < 0.05). Data represent five independent replicates per treatment.

Bio-amendment treatment (5% w/w) produced a dramatic reduction in plasmid transfer efficiency, decreasing HGT frequency to 1.68 ± 0.16 × 10⁻³ (log_10_: −2.77 ± 0.04) with corresponding transconjugant abundance of 1.3 ± 0.2 × 10⁴ CFU g⁻¹ soil. This represented a statistically significant 74.7% reduction in HGT frequency (t = 8.42, p < 0.001), equivalent to a 0.59 log₁₀ unit decrease (Sørensen et al., 2005). The difference in absolute transconjugant numbers was even more pronounced (75.9% reduction), indicating that bio-amendment simultaneously reduced both recipient population density and conjugation efficiency (Zhang et al., 2019). Donor strain (*B. subtilis* 168) survival differed substantially between conditions: control microcosms maintained 42.0% donor recovery (4.2 × 10⁶ CFU g⁻¹), while bio-amended soil reduced donor survival to 28.0% (2.8 × 10⁶ CFU g⁻¹), suggesting the amendment’s antimicrobial activity selectively targeted the Gram-positive donor strain (Heuer et al., 2012). Recipient strain (*E. coli* MG1655, Gram-negative) demonstrated comparable survival between conditions (control: 82.0%, treatment: 79.0%), confirming the amendment’s specificity for Gram-positive bacteria without affecting Gram-negative recipient populations. This differential microbial impact explains the observed HGT reduction: decreased donor density combined with potential biofilm disruption (Fan et al., 2019) and reduced mating pair formation efficiency collectively suppressed *arsR* plasmid dissemination in bio-amended soil. These findings demonstrate that the proposed soil amendment produces dual benefits for resistance gene management: direct suppression of Gram-positive ABR-harbouring populations while simultaneously reducing conjugation-mediated gene transfer efficiency, creating compounded protective effects against antibiotic resistance spread in agricultural systems (Binh et al., 2008).

## Conclusion

This study reveals sublancin 168 as a multifunctional soil amendment addressing climate change-exacerbated threats to agricultural sustainability. Temperature-optimised biofilm formation significantly enhances arsenic sequestration, with 74% increased accumulation at 35°C, providing bioremediation capacity relevant to warming agricultural soils. The selective suppression of Gram-positive bacteria while preserving Gram-negative communities offers ecological balance, preventing pathogenic blooms while maintaining beneficial microbial functions. Most critically, sublancin suppresses antibiotic resistance gene horizontal transfer by 74.7%, operating through dual mechanisms: selective donor strain inhibition and reduced conjugation efficiency. Gene expression analysis reveals coordinated activation of stress-response pathways (arsenic resistance, PTS systems, biofilm formation) while simultaneously suppressing classical antibiotic resistance mechanisms, indicating sophisticated cellular prioritisation favouring antimicrobial production over resistance.

The temperature-dependent nature of sublancin production creates an adaptive advantage under climate change scenarios, with optimal efficacy at 30°C, a temperature projected within decades across major agricultural regions. Integration of thermodynamic, microbiological, and molecular approaches demonstrates that environmental stressors can be harnessed to simultaneously maximise beneficial microbial functions while suppressing pathogenic resistance dissemination. These findings support sublancin’s development as a sustainable, climate-adaptive soil amendment addressing multiple intertwined threats: arsenic contamination, antibiotic resistance spread, and declining soil health. Future work should explore field-scale trials, investigate sublancin’s effects on plant microbiomes, and determine long-term persistence and bioavailability in diverse soil types to translate these laboratory findings into transformative agricultural management strategies under anthropogenic climate change.

## Acknowledgement

This work was funded by the Marie Skłodowska-Curie-UKRI Postdoctoral Fellowship scheme, United Kingdom, with a File number 101152605 and Council reference-EP/Z002664/1. A.M. acknowledge the support from Imperial College London Library during this study and funding for the publication.

## Author’s contribution and declaration

AM: conceptualisation, experiments, analysis, writing & revision; IKL: experiment, writing & revision; MB: revision; TRC: analysis & revision.

The authors declare there is no conflict of interest regarding this work and manuscript.

