## Supplementary files for "Temperature-dependent biofilm and sublancin production immobilise arsenic and antibiotic resistance gene mobility in soil system"

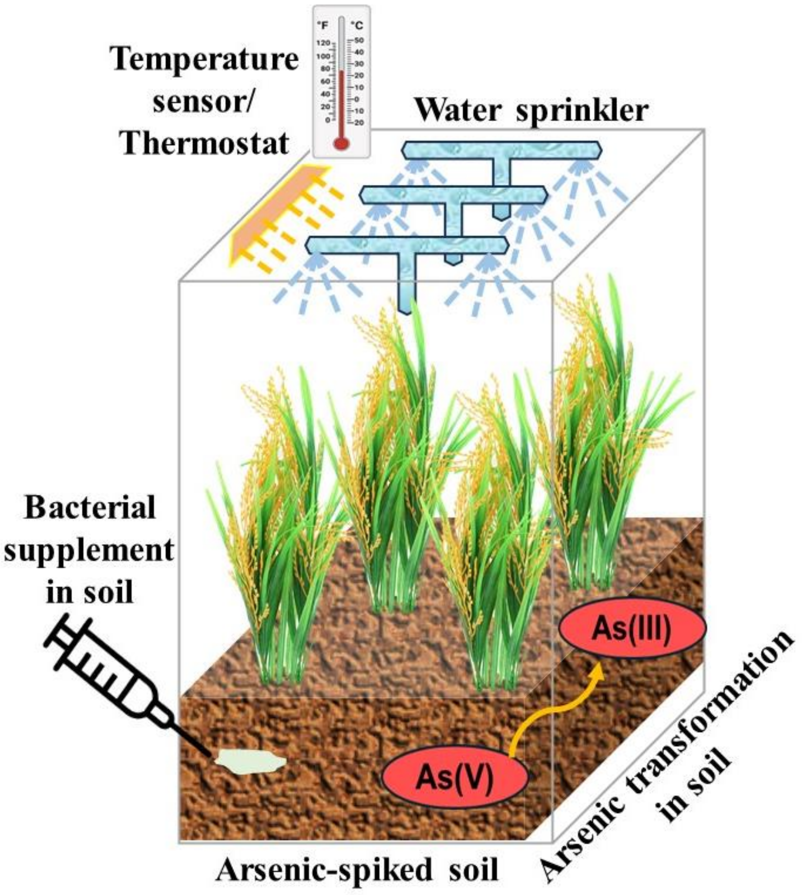


Fig. S1. Experimental setup of the soil microcosm with a thermostat-controlled temperature monitor, As-spiked soil and supplemented bacterium *B. subtilis* 168. Rice plants and water sprinklers were used to keep the soil profile natural, as in fields.

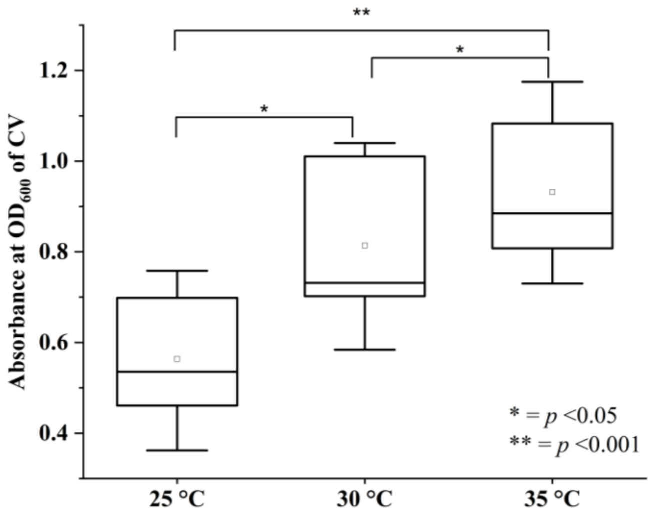


Fig. S2. Total biofilm formation under three different temperature setup. The absorbance of light by crystal violet (CV) at $OD_{600}$ indicates the production of biofilm. Statistical significance was tested at two different levels.

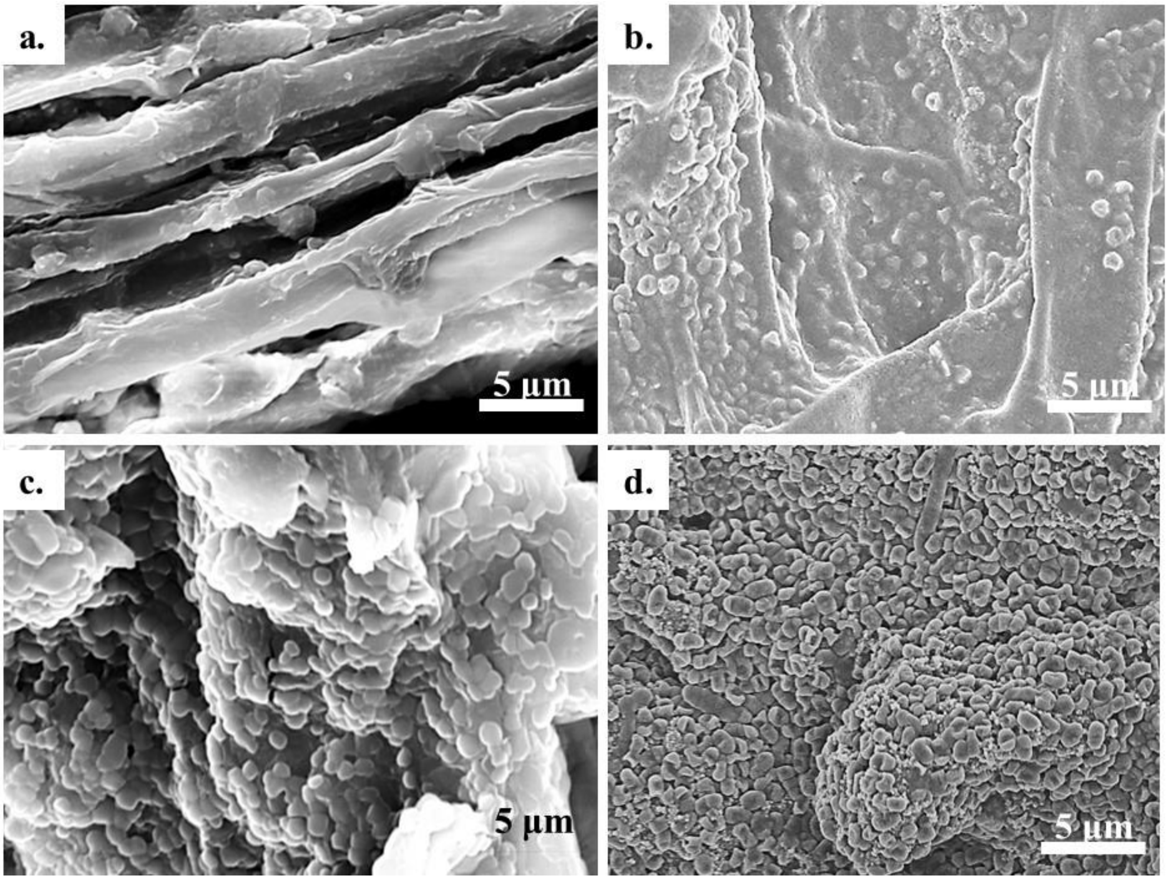


Fig. S3. Temperature-dependent biofilm formation by B. subtilis 168 on the rice plant root surface. Control plant (a) without any bacterial supplementation was compared with the B. subtilis-supplemented plants (b-d), grown under three different temperatures, 25 °C, 30 °C and 35 °C.

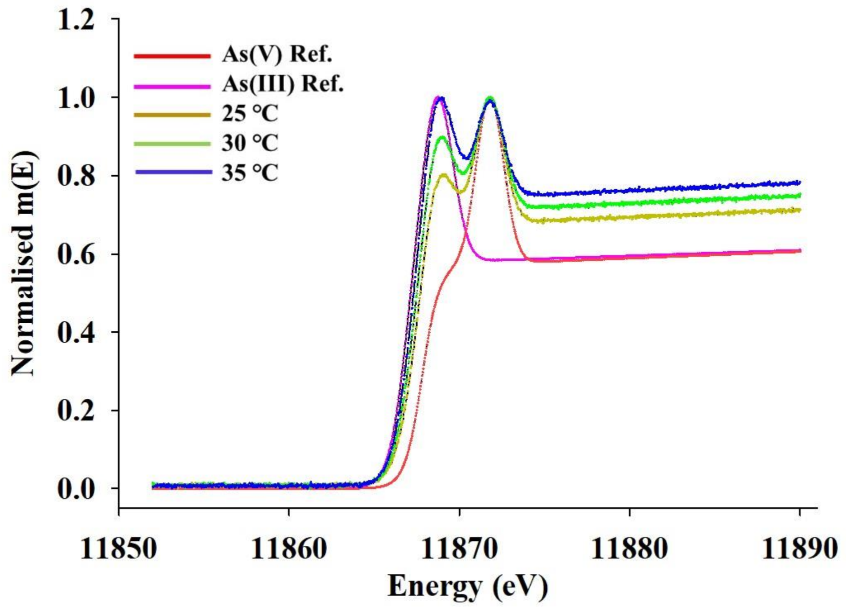


Fig. S4. Arsenic speciation in soil samples with varied temperatures. HPLC-ICP-MS spectra for soil arsenic species were compared with monosodium arsenate and monosodium arsenite as standard reference materials.

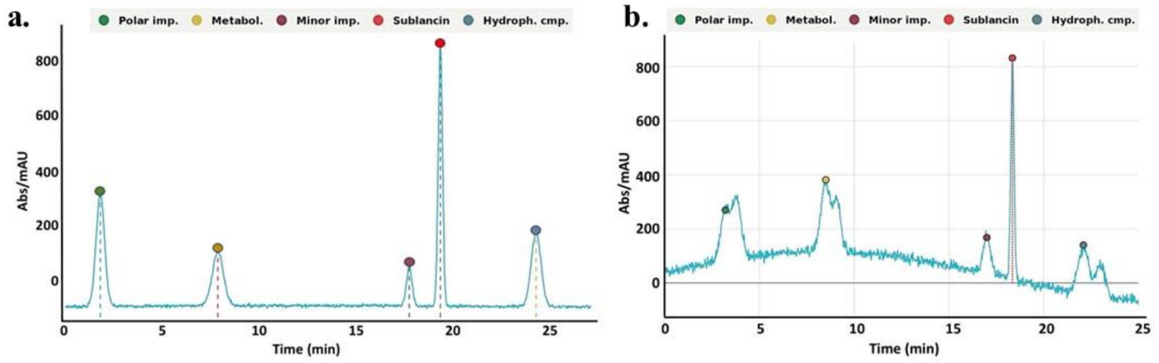


Fig. S5. Extracted sublancin 168 from bacterial growth media (a) and soil samples (b) tested for the right identification using HPLC and further used to test its effect on ABR genes.

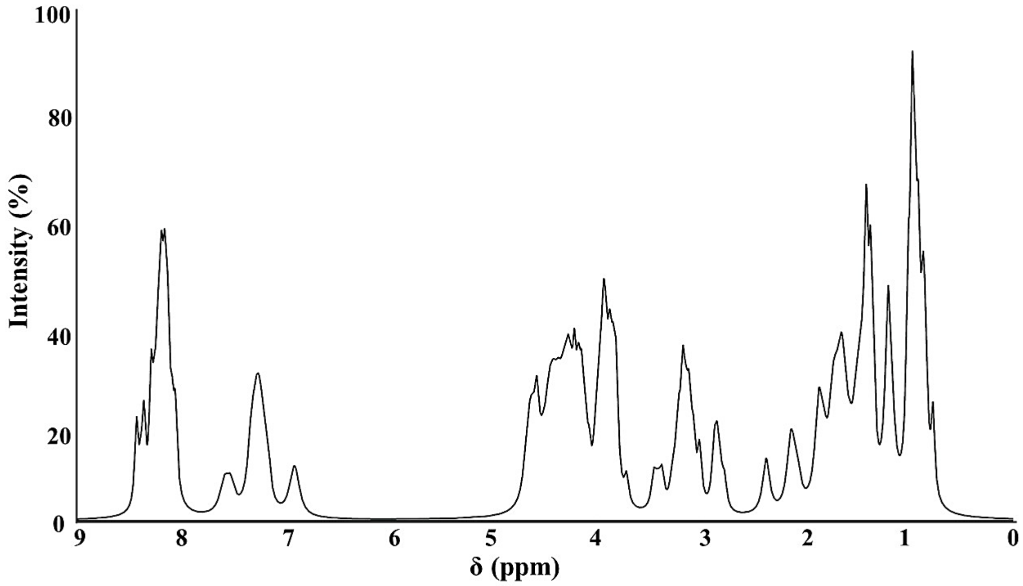


Fig. S6. Sublancin purity analysis using $^{1}$H-NMR. The chemical shift (δ) vs intensity plot is characteristic of a peptide structure with distribution of backbone amide protons, aromatic residues, and diverse aliphatic side chains from the 37-residue sublancin 168 sequence.

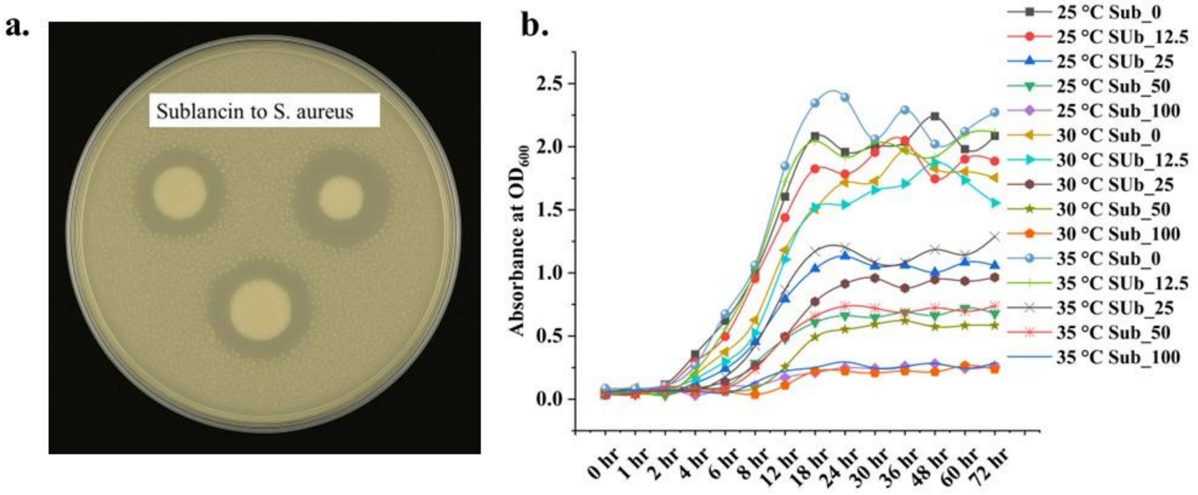


Fig. S7. Effect of sublancin 168 on *Staphylococcus aureus* growth on LB agar plate showing zone of inhibition (a) and in LB broth medium with four sublancin concentrations showing inhibition during growth curve (b).

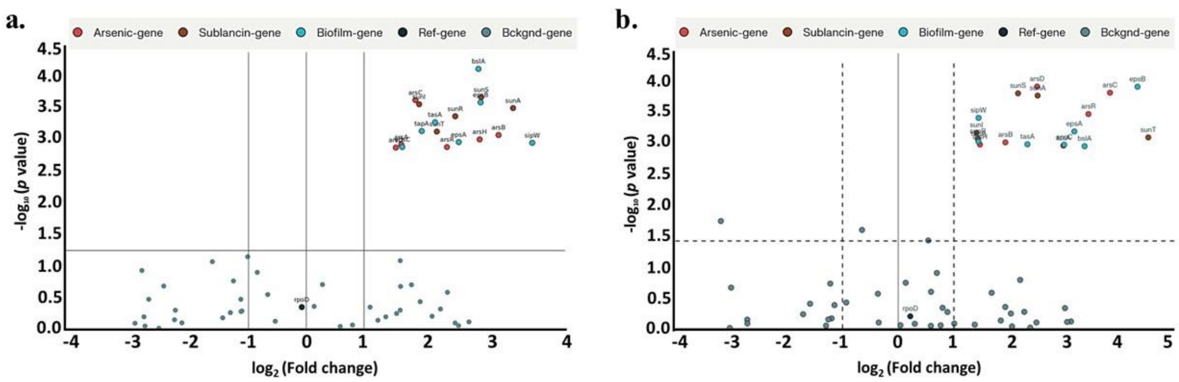


Fig. S8. Scattered volcano plots of As-responsive, sublancin-related and biofilm-forming genes under 25 °C and 35 °C (a-b). Here, the rpoD gene was considered as a reference gene that was unaffected by any temperature change.

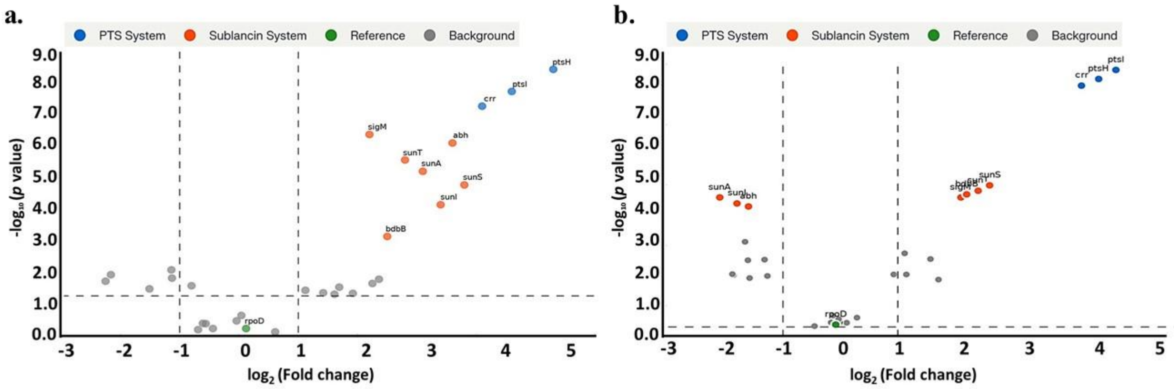


Fig. S9. Scattered volcano plots of sublancin-related and PTS system genes under 25 ℃ and 35 ℃ (a-b). Here, the rpoD gene was considered as a reference gene that was unaffected by any temperature change.

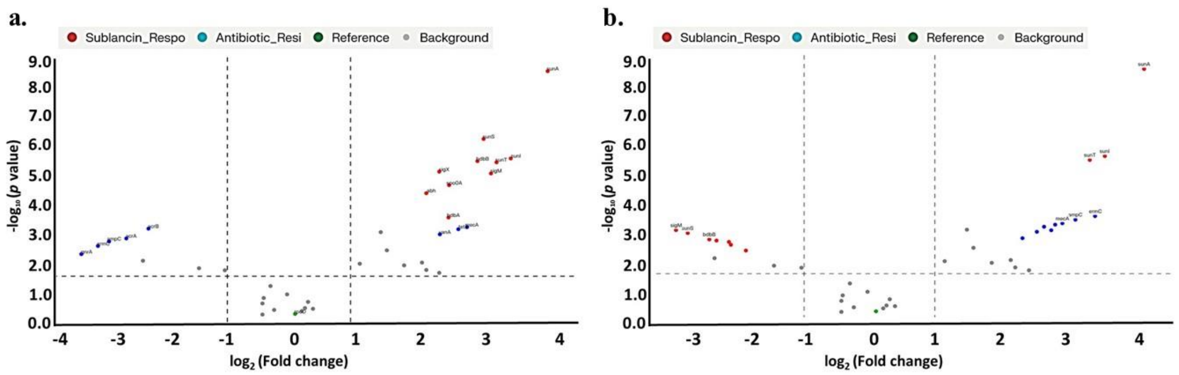


Fig. S10. Scattered volcano plots of sublancin-related and antibiotic-resistance genes under 25 °C and 35 °C (a-b). Here, the rpoD gene was considered as a reference gene that was unaffected by any temperature change.

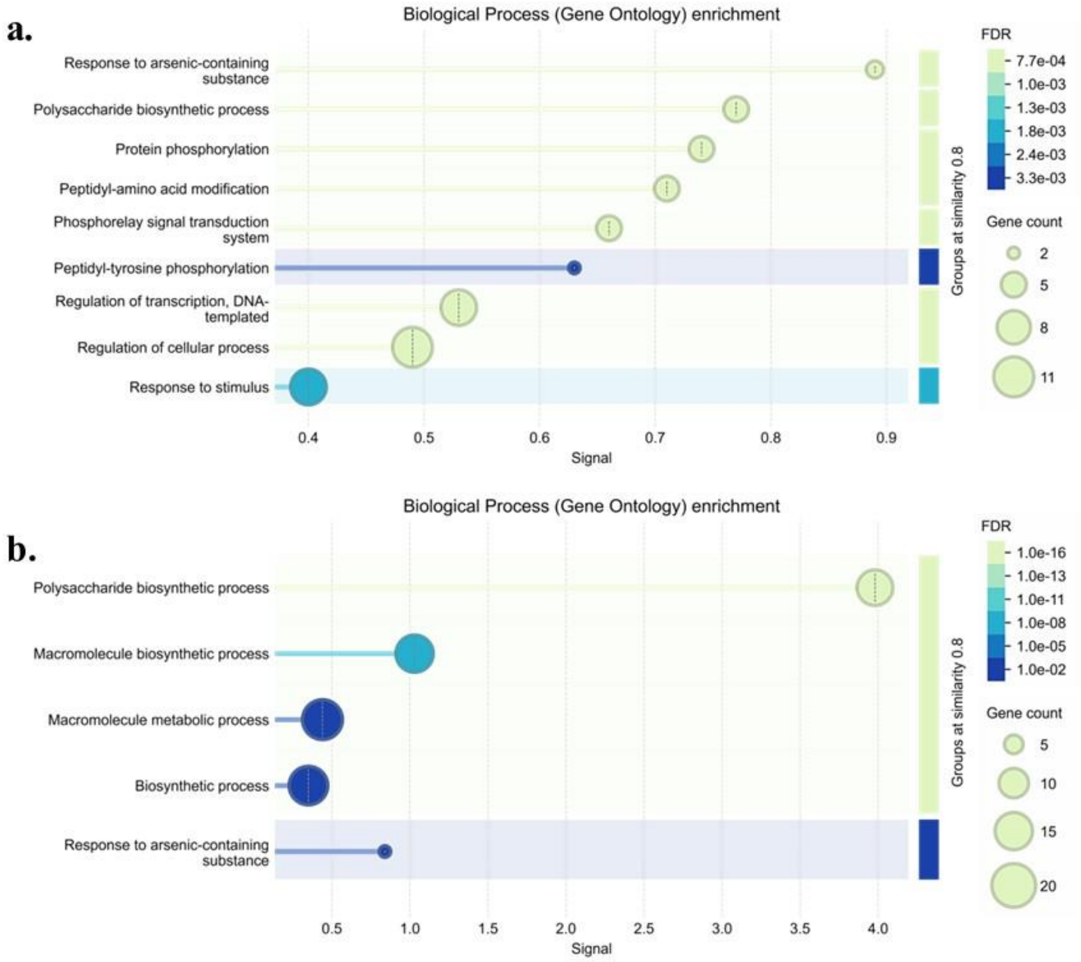


Fig. S11. Gene ontology enrichment and relative fold change of arsenic-responsive (a) and biofilm-producing genes (b) and related cellular activities. Some of the key activities that are directly dependent on these gene functions are shown here.

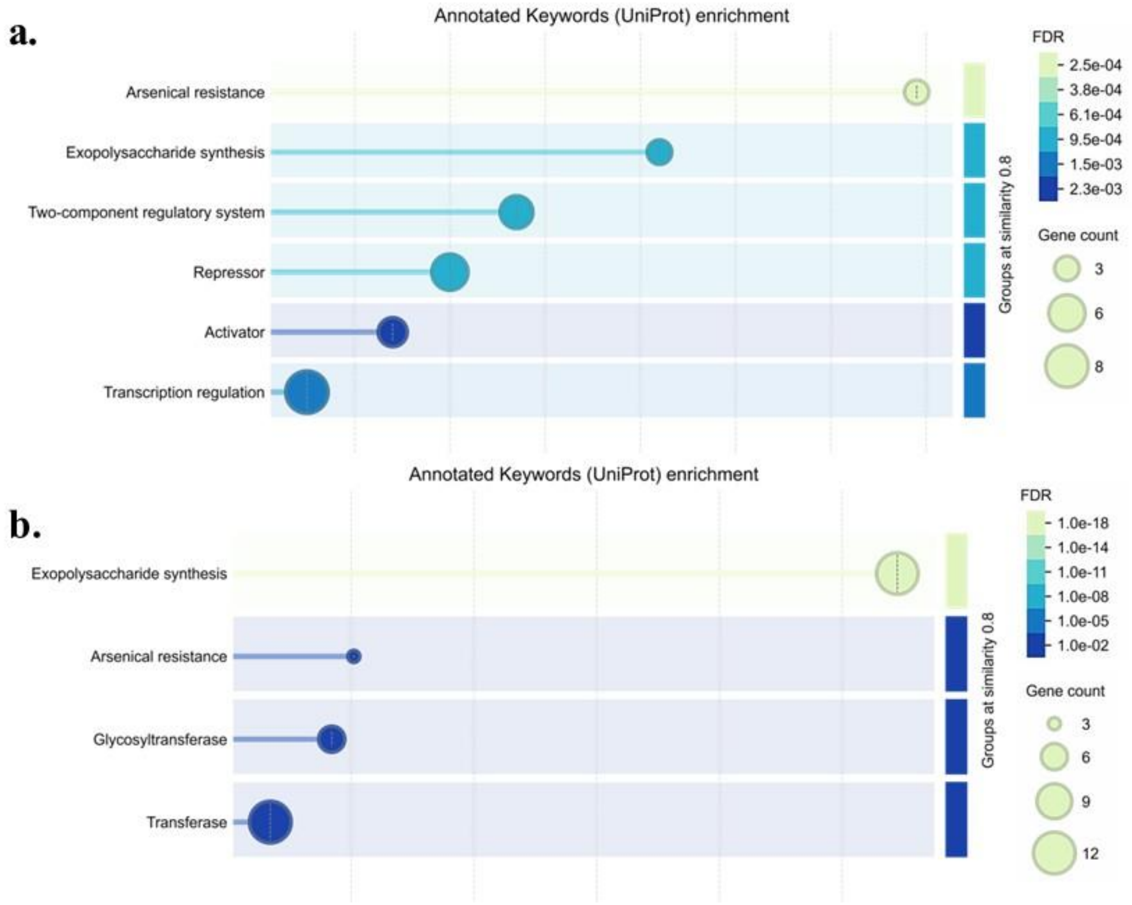


Fig. S12. Relative fold change of genes linked to the UniProt annotations of arsenic-responsive (a) and biofilm-producing (b) molecular activities. These genes are representatives of protein functions as mentioned in the UniProt database.

### Additional discussion on the supplementary figures

#### Arsenic speciation in soil

The HPLC-ICP-MS spectral profiles (Fig. S4) were compared with reference standards of arsenate [As(V)] and arsenite [As(III)]. The As(V) reference spectrum (red) showed a characteristic absorption edge at ~11872 eV, while As(III) (pink) exhibited a distinct shift with reduced post-edge intensity. Soil samples across all three temperatures displayed mixed spectral features, indicating the coexistence of As(V) and As(III) species. At 25 °C (green), As(V) features predominated, with higher post-edge intensity aligning closer to the As(V) standard. At 30 °C (olive), the relative contribution of As(III) increased, evidenced by the partial overlap with the As(III) standard. At 35 °C (blue), spectra displayed a stronger resemblance to As(III), with reduced As(V) peaks and higher absorption intensity consistent with a redox shift toward the more reduced form. Since As(III) is more mobile and toxic than As(V), these findings highlight an important environmental implication: rising soil temperatures could exacerbate arsenic bioavailability and risk in agricultural systems. The results underscore the interplay between climatic factors and arsenic biogeochemistry, suggesting that global warming may amplify arsenic mobilisation and toxicity through temperature-driven redox dynamics in soils.

#### Gene expression in differential conditions

The comparative volcano plot analysis revealed distinct temperature-dependent gene expression patterns between 25°C and 35°C conditions, demonstrating significant differential regulation of arsenic-responsive, sublancin-related, and biofilm formation genes under thermal stress (Fig. S9). The contrasting patterns between the two temperature conditions provide insights into bacterial adaptive strategies and gene regulatory networks responding to environmental temperature fluctuations. At 25°C (panel a), the volcano plot showed moderate gene expression changes with several arsenic-responsive genes (red dots) exhibiting significant upregulation (fold changes +2 to +3 $\log_2$, p-values < 0.001). Key arsenic resistance genes, including arsR, arsB, and arsC, clustered in the upper right quadrant, indicating coordinated activation of metalloid detoxification pathways under cooler conditions (Yang et al., 2012;

Rosen, 2002). Sublancin-related genes (brown dots) demonstrated mixed expression patterns, with some showing moderate upregulation while others remained near baseline, suggesting temperature-dependent regulation of antimicrobial peptide biosynthesis (Wang et al., 2015). Biofilm formation genes (blue dots) exhibited significant upregulation, particularly tasA and epsA, reflecting enhanced extracellular matrix production characteristic of biofilm formation at lower temperatures (Bisht et al., 2023). At 35°C (panel b), dramatic shifts in gene expression patterns emerged, with enhanced separation between significantly upregulated and downregulated genes. Arsenic-responsive genes showed more pronounced upregulation compared to 25°C conditions, with several targets achieving higher statistical significance ($p < 0.0001$) and greater fold changes (+3 to +4 $\log_2$). This enhanced arsenic gene response reflects temperature-dependent amplification of metalloid stress responses, where elevated temperatures may increase arsenic toxicity and require more robust detoxification mechanisms (Li et al., 2022; Nordstrom, 2002). Sublancin genes displayed stronger upregulation at 35°C, with sunA, sunI, and related biosynthesis genes showing significant expression increases, consistent with temperature-dependent optimisation of antimicrobial peptide production (Dorenbos et al., 2002).

The biofilm genes showed interesting temperature-dependent regulation patterns, with some targets maintaining upregulation while others shifted toward downregulation at 35°C. This differential response may reflect temperature-specific biofilm strategies, where higher temperatures favour different biofilm architectures and matrix compositions compared to cooler conditions (Kim et al., 2020; Townsley and Yildiz, 2015). The enhanced scatter and separation observed at 35°C suggest more pronounced stress responses and coordinated gene regulatory networks activated under thermal stress conditions. The reference gene rpoD maintained stable expression across both temperatures, confirming appropriate experimental normalisation and validating the differential expression patterns observed for target genes. The background genes (grey dots) clustered appropriately around the baseline in both conditions, supporting the statistical robustness of the volcano plot analysis. These temperature-dependent expression patterns have significant implications for understanding bacterial adaptation strategies under climate change scenarios. The enhanced gene expression responses at 35°C suggest that warming temperatures may amplify bacterial stress responses, potentially leading to increased arsenic resistance, enhanced antimicrobial production, and altered biofilm formation strategies (MacFadden et al., 2018; Rodríguez-Verdugo et al., 2020). The coordinated upregulation of multiple stress response systems at elevated temperatures indicates synergistic cellular adaptations that may enhance bacterial survival but also contribute to increased antibiotic resistance and altered ecological interactions in warming environments. The comparative volcano plot analysis revealed distinct temperature-dependent expression patterns for sublancin-related and antibiotic resistance genes between 25°C and 35°C conditions, demonstrating contrasting regulatory responses to thermal stress (Fig. S10). The differential gene expression profiles provide insights into bacterial adaptive strategies under varying temperature conditions and highlight the complex interplay between antimicrobial peptide production and resistance mechanisms. At 25°C (panel a), sublancin-related genes (red dots) exhibited significant upregulation with fold changes ranging from +2 to +4 $\log_2$ and high statistical significance ($p < 0.001$). Key sublancin biosynthesis and immunity genes, including sunA, sunI, and sunT, clustered in the upper right quadrant, indicating coordinated activation of antimicrobial peptide production pathways at lower temperatures (Dorenbos et al., 2002; Wang et al., 2015). This enhanced sublancin gene expression is consistent with established temperature-dependent regulation where cooler conditions favour sublancin biosynthesis, potentially as an adaptive strategy for competitive advantage in environmental conditions typical of soil ecosystems (Ji et al., 2015).

Antibiotic resistance genes (blue dots) showed moderate expression at 25°C, with several ARGs displaying significant upregulation (fold changes +1.5 to +3 $\log_2$, $p < 0.01$). The distribution of resistance genes suggests baseline ARG expression maintained under standard growth conditions, reflecting constitutive resistance mechanisms that provide broad-spectrum protection against various antimicrobial compounds (Andersson et al., 2016; Forsberg et al., 2012). At 35°C (panel b), a dramatic shift in expression patterns emerged, with sublancin genes showing reduced upregulation compared to 25°C conditions, while antibiotic resistance genes demonstrated enhanced expression. Several sublancin-related targets exhibited lower fold changes (+1 to +2 $\log_2$) and reduced statistical significance, suggesting temperature-dependent suppression of antimicrobial peptide production at elevated temperatures. This pattern aligns with previous studies demonstrating that sublancin stability and activity decrease at higher temperatures, potentially reducing the selective pressure for sublancin gene expression (Ji et al., 2015; Wang et al., 2015). In contrast, antibiotic resistance genes showed enhanced upregulation at 35°C, with multiple ARGs achieving higher fold changes (+2 to +4 $\log_2$) and greater statistical significance compared to 25°C conditions. This temperature-dependent ARG enhancement reflects established relationships between elevated temperatures and increased antibiotic resistance gene expression, mediated through heat-shock response networks and enhanced horizontal gene transfer efficiency (MacFadden et al., 2018; Li et al., 2022). The enhanced ARG expression may represent compensatory mechanisms where bacteria prioritise conventional resistance systems when antimicrobial peptide production becomes less favourable under thermal stress. The inverse relationship between sublancin and antibiotic resistance gene expression at different temperatures suggests temperature-dependent trade-offs in bacterial defence strategies. At 25°C, the predominance of sublancin gene upregulation indicates prioritisation of offensive antimicrobial production, while at 35°C, the shift toward antibiotic resistance gene expression suggests defensive resistance mechanisms become more favourable under thermal stress conditions (Spohn et al., 2019). These temperature-dependent patterns have significant implications for understanding microbial community dynamics under climate change scenarios. The reduced sublancin production at elevated temperatures, combined with enhanced antibiotic resistance gene expression, suggests that global warming may shift bacterial competitive strategies away from antimicrobial peptide-mediated competition toward conventional antibiotic resistance-based survival, potentially altering soil microbial ecology and contributing to environmental resistance reservoirs (Rodríguez-Verdugo et al., 2020).

The comparative volcano plot analysis revealed striking temperature-dependent coordination between the PTS system and sublancin-related gene expression, demonstrating remarkable synchronisation between these functionally interconnected cellular systems at different temperatures (Fig. S11). The expression patterns provide molecular evidence for the established functional dependency between sublancin antimicrobial activity and PTS system components, while revealing temperature-specific regulatory dynamics that govern their co-expression. At 25°C (panel a), both the PTS system genes (blue dots) and the sublancin system genes (red dots) exhibited pronounced upregulation with exceptional statistical significance. PTS genes, including ptsI, ptsH, and crr, clustered in the upper right quadrant with fold changes ranging from +3 to +4 $\log_2$ and p-values exceeding 8.0 ($-\log_{10}$). Similarly, sublancin genes showed coordinated upregulation with sunA, sunI, and sunT, achieving fold changes of +2 to +3 $\log_2$ and comparable statistical significance (Garcia De Gonzalo et al., 2011; Oman et al., 2015). This synchronised expression pattern reflects the obligate functional relationship where sublancin requires active PTS components for its antimicrobial mechanism, and bacterial sensitivity to sublancin depends on functional phosphotransferase activity (Biswas et al., 2021). At 35°C (panel b), the coordinated expression patterns were maintained but with modified intensity profiles. PTS system genes continued to show significant upregulation with ptsI and ptsH maintaining high statistical significance ($p < 0.001$) and substantial fold changes (+3 to +4 $\log_2$). Sublancin genes displayed somewhat reduced expression levels compared to 25°C conditions, with most targets showing fold changes of +1.5 to +2.5 $\log_2$ but retaining high statistical significance. This temperature-dependent modulation may reflect thermal stability considerations for sublancin peptide, where elevated temperatures could affect peptide folding, stability, or activity, leading to adjusted biosynthetic gene expression (Wang et al., 2015; Ji et al., 2015).

The consistent co-upregulation of PTS and sublancin genes across both temperature conditions provides strong molecular support for their mechanistic interdependence. Previous studies have demonstrated that sublancin's antimicrobial activity requires functional PTS components, particularly HPr (ptsH) and PtsG, with mutations in these genes conferring sublancin resistance (Oman et al., 2015). The coordinated gene expression patterns observed here suggest that bacterial cells enhance both systems simultaneously, potentially to maximise sublancin production efficiency while ensuring adequate PTS machinery for the peptide's mechanism of action. The functional relationship manifests through sublancin's unique mechanism, where the antimicrobial peptide requires PTS-mediated processes for cellular entry, phosphorylation, or target interaction (Biswas et al., 2021). The S-linked glucose modification on sublancin may be recognised by PTS transporters, facilitating peptide entry into target cells where PTS-mediated phosphorylation could activate sublancin's bactericidal properties (Garcia De Gonzalo et al., 2011). The coordinated upregulation ensures that sublancin-producing bacteria maintain sufficient PTS activity to support optimal antimicrobial peptide function. The reference gene rpoD maintained stable expression across both conditions, validating the experimental approach and confirming that the observed changes represent genuine temperature-dependent regulation rather than technical artefacts. Background genes clustered appropriately around baseline levels, supporting the statistical robustness of the differential expression analysis. These results have significant implications for understanding sublancin's ecological role and potential applications. The temperature-dependent coordination between PTS and sublancin systems suggests that environmental temperature fluctuations could modulate antimicrobial peptide efficacy by affecting both biosynthetic capacity and functional machinery. The synchronised expression patterns also provide molecular targets for enhancing sublancin production in biotechnological applications, where co-optimisation of both PTS and sublancin genes could improve antimicrobial peptide yields and therapeutic potency.

**Gene fold change and enrichment terms**

Gene Ontology (GO) enrichment analysis revealed distinct biological process profiles between the two experimental conditions, with markedly different significance levels and functional categories being enriched (Fig. S12). The comparative analysis demonstrates condition-specific cellular responses with varying degrees of statistical support and gene representation across multiple biological pathways. In the first condition (panel a), arsenic-responsive processes dominated the enrichment profile, with "Response to arsenic-containing substance" achieving the highest statistical significance (FDR ~7.7e-04) and moderate gene representation (~8 genes). This category reflects activation of metalloid detoxification pathways essential for bacterial survival under heavy metal stress (Rosen, 2002; Yang et al., 2012). Polysaccharide biosynthetic processes showed significant enrichment (FDR ~1.0e-03), indicating enhanced extracellular matrix production likely associated with biofilm formation and cellular protection mechanisms under stress conditions (Rehm, 2010; Schmid et al., 2015). Post-translational modification pathways were prominently enriched, including "Protein phosphorylation," "Peptidyl-amino acid modification," and "Peptidyl-tyrosine phosphorylation" categories with FDR values ranging from 1.3e-03 to 2.4e-03. These enrichments suggest enhanced regulatory networks involving protein modifications essential for stress responses and cellular adaptation (Hunter, 2000; Stock et al., 2000). The "Phosphorelay signal transduction system" enrichment indicates activation of two-component regulatory systems critical for environmental sensing and adaptive responses (Capra and Laub, 2012). In the second condition (panel b), dramatically enhanced enrichment patterns emerged with much stronger statistical significance across multiple categories. "Response to arsenic-containing substance" maintained high significance but with increased signal strength (~2.5 vs ~0.9), suggesting amplified arsenic stress responses under this experimental condition. "Polysaccharide biosynthetic process" showed exceptional statistical significance (FDR ~1.0e-16) and substantial gene representation (~15 genes), indicating massive upregulation of extracellular matrix biosynthesis.

Most notably, broad biosynthetic and metabolic categories achieved extraordinary enrichment levels in condition b. "Macromolecule biosynthetic process" and "Macromolecule metabolic process" displayed FDR values of 1.0e-11 and 1.0e-05 respectively, with large gene counts (15-20 genes each). These categories represent fundamental cellular processes including protein synthesis, nucleic acid metabolism, and complex molecule biosynthesis, suggesting comprehensive metabolic reprogramming under the second experimental condition (Ralser, 2018). The "Biosynthetic process" category showed exceptional statistical support (FDR ~1.0e-02) with the highest gene representation (~20 genes), indicating coordinated activation of anabolic pathways. This pattern suggests that the second condition promotes extensive biosynthetic activity, potentially reflecting enhanced growth conditions, stress-induced metabolic shifts, or specific environmental stimuli that favour increased cellular biosynthesis (Locasale and Cantley, 2011). The contrasting enrichment profiles between conditions indicate fundamentally different cellular states and adaptive strategies. Condition a appears to represent moderate stress responses with specific activation of arsenic detoxification and regulatory pathways, while condition b demonstrates extensive metabolic activation with enhanced biosynthetic capacity and amplified stress responses. These patterns suggest that condition b may involve more favourable growth conditions or specific stimuli that promote comprehensive cellular activation beyond simple stress responses.

UniProt annotated keyword enrichment analysis revealed distinct functional category profiles between the two experimental conditions, demonstrating condition-specific cellular responses with varying statistical significance levels and gene representation patterns (Fig. S13). The comparative analysis highlights fundamental differences in cellular priorities and metabolic adaptations under the respective experimental treatments. In the first condition (panel a), arsenical resistance emerged as the most statistically significant category (FDR ~2.5e-04) with moderate gene representation (~3 genes), indicating specific activation of heavy metal detoxification pathways essential for bacterial survival under metalloid stress (Rosen, 2002; Chen et al., 2016). Exopolysaccharide synthesis showed strong enrichment (FDR ~3.8e-04) with substantial gene count (~8 genes), reflecting enhanced extracellular matrix production characteristic of biofilm formation and protective responses to environmental stress (Flemming and Wingender, 2010; Rehm, 2010). Regulatory system categories were prominently enriched, including "Two-component regulatory system" (FDR ~6.1e-04, ~6 genes) and "Transcription regulation" (FDR ~1.5e-03, ~8 genes). These enrichments indicate activation of sophisticated regulatory networks involving environmental sensing, signal transduction, and coordinated gene expression responses critical for bacterial adaptation to changing conditions (Capra and Laub, 2012; Gao and Stock, 2009). The presence of both "Repressor" and "Activator" categories suggests complex transcriptional control mechanisms where both positive and negative regulation coordinate cellular responses to experimental conditions.

In the second condition (panel b), exopolysaccharide synthesis achieved exceptional statistical significance (FDR ~1.0e-18) with increased gene representation (~9 genes), indicating massive upregulation of extracellular matrix biosynthesis. This dramatic enhancement compared to condition a suggests that condition b provides more favourable conditions or specific stimuli that promote extensive biofilm matrix production (Branda et al., 2005; Vlamakis et al., 2013). Arsenical resistance maintained significant enrichment (FDR ~1.0e-05) but with reduced relative prominence compared to condition a, suggesting that while heavy metal stress responses remain active, other metabolic processes become prioritised under condition b. The emergence of glycosyltransferase and transferase categories with high statistical significance (FDR ~1.0e-05 to 1.0e-02) indicates enhanced enzymatic activity related to carbohydrate modification and metabolic processing (Lairson et al., 2008; Campbell et al., 1997). The transferase category achieved the highest gene count (~12 genes) in condition b, representing broad enzymatic activities including phosphoryl transfer, glycosyl transfer, and other metabolic transformations. This extensive enrichment suggests comprehensive metabolic activation where multiple transferase-mediated pathways become coordinately upregulated, potentially reflecting enhanced biosynthetic capacity or specific metabolic requirements under condition b (Copley and Bork, 2000). The contrasting enrichment profiles reveal fundamentally different cellular strategies between conditions. Condition a appears to emphasise stress response and regulatory control, with moderate exopolysaccharide production and specific arsenical resistance mechanisms. Condition b demonstrates extensive metabolic activation with dramatically enhanced exopolysaccharide synthesis and broad enzymatic pathway engagement, suggesting more favourable growth conditions or specific environmental stimuli that promote comprehensive cellular biosynthesis. These patterns have important implications for understanding bacterial adaptation mechanisms under varying environmental conditions. The condition-dependent shifts from stress-focused responses (condition a) to biosynthesis-oriented metabolism (condition b) suggest that bacterial communities can dynamically adjust their functional priorities based on environmental cues and resource availability (Ralser, 2018; Locasale and Cantley, 2011).
